# Dendritic computation for rule-based flexible categorization

**DOI:** 10.1101/2025.06.03.657766

**Authors:** Yuan Zhang, Yanhe Liu, Yachuang Hu, Zhenkun Zhang, Jie Du, Lele Cui, Qi Liu, Lin Zhong, Yu Xin, Ruiming Chai, Li Deng, Jingwei Pan, Jackie Schiller, Ning-long Xu

**Affiliations:** Institute of Neuroscience, State Key Laboratory of Brain Cognition and Brain-inspired Intelligence Technology, Center for Excellence in Brain Science and Intelligence Technology, Chinese Academy of Sciences, Shanghai 200031, China; University of Chinese Academy of Sciences, Beijing 100049, China; Shanghai Center for Brain Science and Brain-Inspired Intelligence Technology, Shanghai 201210, China; Department of Physiology, Technion Medical School, Bat-Galim, Haifa 31096, Israel

## Abstract

A hallmark of intelligent behavior is the ability to flexibly respond to external sensory inputs based on dynamically changing rules. A central question is how neurons in the brain implement computations underlying intelligent behaviors. The neocortical pyramidal neurons use their elaborated dendritic arbors to segregate a plethora of inputs and dynamically integrate them—a process known as dendritic computation—which may play important roles in rule-dependent sensory processing. However, evidence directly linking dendritic computation with intelligent cognitive behaviors has been absent. Here we combine two-photon imaging and a rule-switching flexible categorization task in mice to show that a projectome-defined extratelencephalic (ET) cortical layer 5 (L5) neurons in the auditory cortex integrate dendritic rule information and somatic sensory input to enable rule-dependent flexible categorization. The apical dendrite and soma within the same ET neurons exhibit distinct compartmental representations for sensory and rule information, with the soma predominantly encoding sensory information and the dendrites representing inferred task rule. Simultaneous optogenetic dendritic inhibition and two-photon imaging revealed that dendritic rule coding is essential for somatic output of flexible categorization. Our findings indicate that nonlinear dendritic integration of rule and sensory information constitutes a neuronal computational mechanism underlying rule-switching flexible decision-making.

## Introduction

Cortical pyramidal neurons serve as crucial computational units in the brain^1–3^. Almost all of them extend large apical dendrites towards the superficial layer of the cortex, where they branch out extensively to form a tuft tree receiving a wide range of contextual input information ^1,4,5^. Pyramidal neuron dendrites play a crucial role in integrating diverse network inputs and transforming them into spiking outputs that influence perception and behavior^1,4–7^. Since the active role of cortical pyramidal-neuron dendrites was initially proposed^8^, the importance of dendritic processing has been widely demonstrated both in vitro and in vivo during sensory and motor behaviors^9–25^. However, the contributions of dendritic processing to circuit computations underlying cognitive functions, such as decision-making and cognitive flexibility, remain unclear.

Decision-making involves transforming sensory inputs into appropriate motor outputs in a context- and state-dependent manner, where the same sensory input may be mapped to different motor outputs depending on the environmental state or task rule. Although substantial advances have been made in exploring the neural mechanisms underlying decision-making^26–28^, prior research examined neural representations primarily based on point-process neurons, often disregarding the contributions of dendritic mechanisms. Whether and how the dendrites integrate and transform diverse sources of information from various neuronal populations in multiple brain regions during decision-making remains unknown. Consequently, major knowledge gaps persist in our understanding of the neural mechanisms underlying cognition.

Two unique properties of the pyramidal neuron dendrites can be important in information processing during a decision-making task: one is the compartmentalization, arising from the intricate morphology, the biophysical properties of dendritic trees, and the input distribution and statistics^6,12,13,24,29–33^. This property allows for distinct types of information to be represented in different spatial domains within a single neuron, enabling local dendritic operations, parallel processing, and learning for various types of inputs^13,29,32,34,35^. The second property involves the interaction and communication between compartments via active and nonlinear integrative mechanisms of the dendrite^6,7,11,12,14,20,30^, allowing for rich and dynamic input-output transformations.

A previous study^20^ provided evidence for *in vivo* nonlinear dendritic integration during an active perceptual behavior by demonstrating that global dendritic activity associated with Ca^2+^ plateau potentials arises from nonlinear integration of sensory and motor information originating from distinct brain areas. Yet, two important questions remain to be addressed in order to gain insights into dendritic computations underlying cognitive functions. First, while the global dendritic spikes represent the final output of information integration, it remains unclear how task-relevant cognitive and behavioral variables are compartmentalized in dendritic domains. Second, how compartmental representations in different sub-neuronal domains interact during cognitive tasks to produce behaviorally relevant output remains unknown. Addressing these questions has been difficult due to challenges in isolating cognitive processes from complex behavioral outputs and identifying their relationships with dendritic specific activities.

To tackle these questions, we measured sub-neuronal dendritic and somatic activity using *in vivo* two-photon imaging from projection-defined ET L5 pyramidal neurons in auditory cortex (ACx) in mice performing a rule-switching flexible categorization (RSFC) task. The RSFC task was developed in our previous study^36^, which is analogous to the Wisconsin Card Sorting Test (WCST) where subjects infer changing rules through feedback from choice outcomes, and apply these rules to adapt subsequent choices. In the RSFC task, mice are required to categorize various pure tones as ‘high’ or ‘low’ using a predefined classification boundary, i.e., categorization rule. The categorization rule changes randomly in different blocks of trials for multiple times within a behavioral session, where mice need to infer the rule change through reward outcome. We isolated and quantified key cognitive and behavioral variables from the task and identified the underlying computational algorithm. Our results show that the apical dendrites and somas of a projectome-defined ET L5 neurons in ACx exhibit sub-neuronal level compartmental representations for distinct task variables, including task rule, sensory feature, and categorical choices. Task rule information is largely confined to the apical dendritic compartments, whereas sensory information is mainly compartmentalized in the somatic domains. Meanwhile, the category information, as reflected by choices, resulting from the computation that integrates task rule and sensory stimuli, is ubiquitously represented in both dendritic and somatic activity. Furthermore, by combining simultaneous optogenetic dendritic inhibition and two-photon Ca^2+^ imaging, we demonstrate the crucial contribution of dendritic activity to the somatic encoding of rule-dependent category choices. Our findings uncover a single-neuron level computation based on dendritic compartmentalization and nonlinear integration contributing to the cognitive processing of flexible decision-making.

## Results

### Imaging dendritic activity during flexible decision-making behavior

In a dynamic world, rules for sensory-choice-outcome contingencies often change depending on the environmental state. The most efficient way for a decision maker to determine the proper choice is to first estimate the hidden task state and then choose appropriate actions upon given sensory stimuli (Fig. 1a). This requires the integration of sensory information with the estimated task rule information to produce proper choice output. Such information integration could in principle be efficiently carried out by the dendritic mechanisms in cortical pyramidal neurons^7,9,10,14,20^ (Fig. 1i).

**Fig. 1.**
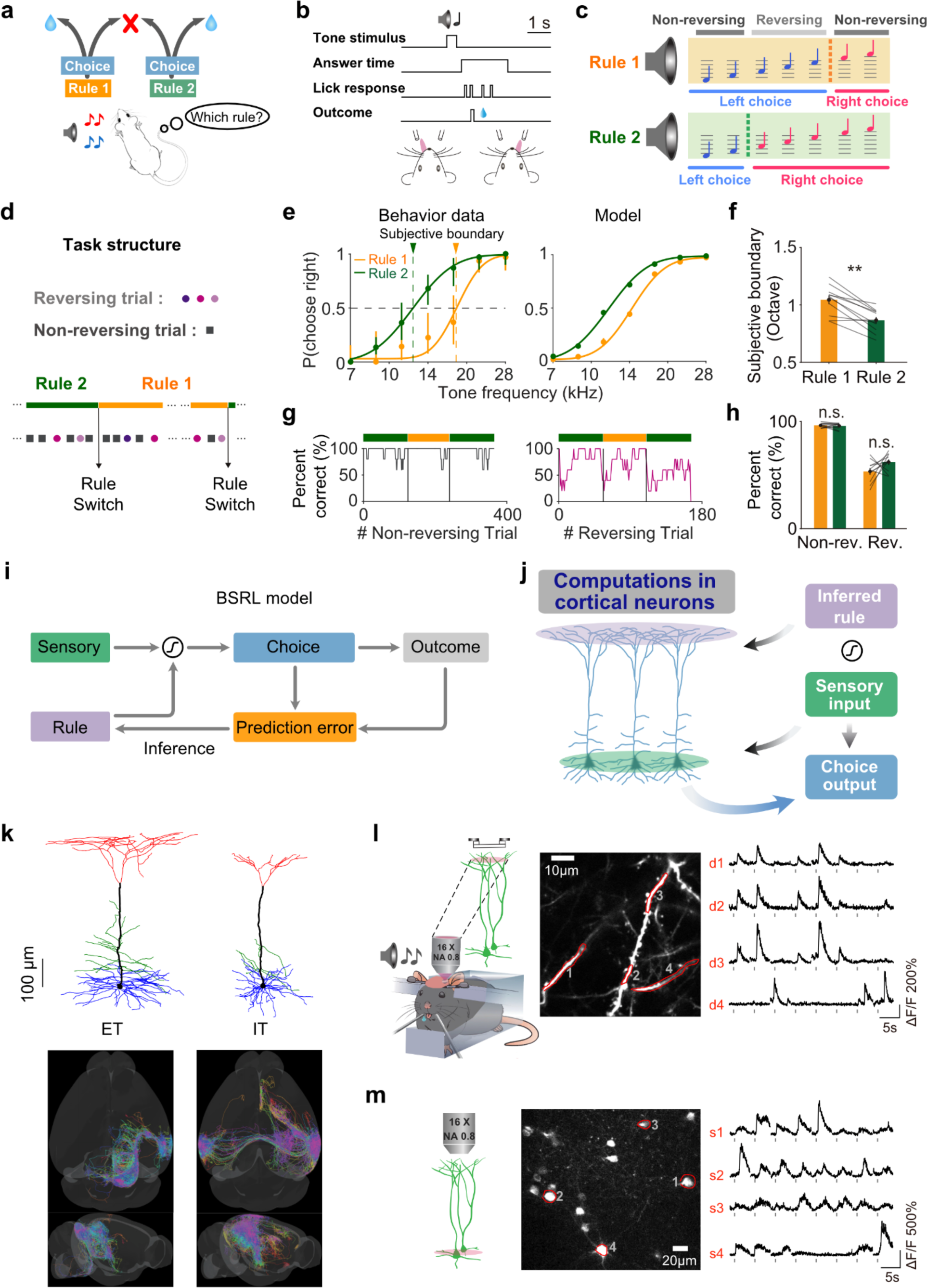
Two-photon imaging of dendrites and soma in projection-defined L5 pyramidal cell types during rule-switching flexible categorization. **a**, Illustration of rule-switching flexible categorization. For the same tone stimulus, mice need to make the choice category based on the rule to get a water reward. **b**, Temporal structure for behavioral events in a single trial. **c**, Schematic showing the auditory-guided rule-switching flexible categorization task with changing boundaries under 2 rules. Yellow, rule 1. Green, rule 2. Dashed line, boundary. **d,** Task structure of example session, reversing and non-reversing trials were randomly given under each rule, every day both rules were encountered by the animal. **e**, Left, psychometric functions from trials under the two categorization rules from an example behavior session. The shift in the subjective category boundary corresponds to the rule switching. Error bars represent 95% confidence intervals. Yellow, from trials under rule 1; green, rule 2. Right, psychometric functions of choice probability obtained by fitting a BSRL model as in **i** to the behavioral data. Error bars represent the SEM. **f**, Comparison of subjective boundaries under different rules. Wilcoxon signed-rank test, p = 0.002, n = 11 mice. Error bars indicate SEM. **g**, Performance of non-reversing trials and reversing trials from an example session under different rules, non-reversing trials stay high correct rate (left), and reversing trials gradually increase the correct rate after each rule transition (right). **h**, Comparison of correct rate between different rules for reversing and non-reversing trials. Wilcoxon signed-rank test, n.s., not significant, p = 0.76 for non-reversing trials, p = 0.15 for reversing trials, n = 11 mice. **i**, Schematic showing the BSRL model for capturing rule-switching flexible categorization behavior based on rule inference (see Methods). **j**, Hypothetical neuronal computation that could implement a categorization rule-dependent perceptual decision-making process. L5 pyramidal neurons in the sensory cortex combine feedback inputs representing internal state (e.g. rule estimation) received at the distal dendritic tuft with feed-forward sensory inputs received at the perisomatic region and produce enhanced output to influence behavioral choices. **k**, Reconstruction of dendrites and axons of single ET and IT neurons in ACx (Methods). Top, example dendritic morphologies of ET and IT neurons. Red, tuft dendrites; green, oblique dendrites; blue, basal dendrites. Bottom, projections of example ET (n = 23) and IT neurons (n = 39). **l**, Two-photon Ca^2+^ imaging from the tuft dendrites of L5 neurons in the ACx expressing GCaMP6s during behavior. Right, example Ca^2+^ imaging traces from indicated dendritic ROIs. Gray rectangle marks indicate sound stimulus time. **m**, Ca^2+^ imaging traces from example somatic ROIs as in **l**.

To examine dendritic computations for complex and flexible behaviors, we trained head-fixed mice to perform an auditory-guided RSFC task^36^ (Fig. 1b-f; see Methods). On a given trial, a head-fixed mouse is presented with a tone stimulus randomly selected from a set of 7 frequencies. The animal needs to determine whether the tone belongs to the ‘high’ or ‘low’ frequency category based on an experimentally defined category boundary and report their choices by licking the left or right lick spout to obtain water reward. The category boundary randomly switches between a lower and higher frequency across different blocks of trials (100-200 trials/block). The tone frequencies between the two category boundaries are categorized as ‘low’ (licking left) when the boundary shifts to the higher frequency (rule 1). Conversely, the same set of tones needs to be categorized as ‘high’ (licking right) when the boundary shifts to the lower frequency (rule 2). Since choices for the tones between the two category boundaries are reversed under the two rules, these tones are termed ‘reversing’ stimuli. Meanwhile, the tones with frequencies higher or lower than the two boundaries (termed ‘non-reversing’ stimuli) remain in the ‘high’ or ‘low’ category without requiring a change in choice upon rule switching. There are no explicit cues for rule switching, meaning that the categorization rule could only be inferred from the choice outcome in reversing trials. The choice behavior with multiple tone frequencies allows us to construct a psychometric function to estimate the point of subjective equality (PSE), which represents the subjective categorization boundary (Fig. 1e)^36–38^. While the degree of categorization (the sharpness of the psychometric curves) could vary across blocks of trials, mice exhibited significant shift of PSE upon rule changes within each session, indicating successful rule switching (Fig. 1f). There were no significant systematic differences in the correct rates for the reversing and non-reversing stimuli between the two rules (Fig. 1h). As described previously^36^, mice are able to learn the task structure to perform rule switching multiple times within a single session (Fig. 1g). The cognitive process of rule inference can be captured and quantified using a belief-state reinforcement learning (BSRL) model (Fig. 1e,i; see Methods)^36^, which allows for quantification of unobservable cognitive variables, including the subjective rule estimate and the value prediction error. This task also contains rich trial types that allow us to dissociate multiple behavioral and cognitive variables. In the non-reversing trials, while the stimulus-choice contingency remains constant over the entire session, the rule information changes across different blocks, allowing the rule-related information to be dissociated from the sensory and choice information. In the reversing trials, when the rule switches, the same sensory stimuli map to different choices; hence, the choice and sensory information can be dissociated.

Different subtypes of L5 cortical pyramidal neurons exhibit distinct structural and functional properties^5,24,39,40^. To examine the dendritic computations in flexible decision-making, we first sought to determine which subtype of cortical L5 pyramidal neurons in ACx is likely involved in our auditory decision-making task. We began by studying the differential morphological and projection patterns of different subtypes of cortical L5 pyramidal neurons. The contralateral ACx-projecting L5 neurons represent a major proportion of intratelencephalic (IT) neurons, while the superior colliculus (SC)-projecting neurons represent a major part of the ET L5 pyramidal neurons^41^ (Extended Data Fig. 1a-c). We used our single-neuron projectome pipeline^42,43^ to reconstruct both the dendritic morphologies and the whole-brain axonal projections of ACx neurons. We obtained 71 contralateral ACx-projecting L5 IT neurons and 39 SC-projecting L5 ET neurons, both with axons and dendrites fully reconstructed for morphological comparison (Fig. 1j; Extended Data Fig. 1d-f). Consistent with prior notions^1,39,44^, the apical dendritic tufts of ET neurons are more widely spread than those of IT neurons and may be able to integrate a broader range of behavior-related information (Fig. 1k; Extended Data Fig. 1g,h). The L5 IT neurons primarily send axons to the contralateral auditory cortex and other sensory and association cortices (Extended Data Fig. 1d,f), whereas the L5 ET neurons send axons to many subcortical regions extensively (Extended Data Fig. 1d,e). These distinct projection patterns suggest that the L5 IT and ET neurons carry out distinct functions: the L5 IT neurons may be involved in coordinating sensory representations across hemispheres and presumably also across different sensory modalities, while the L5 ET neurons are likely responsible for transforming auditory representations into behavioral outputs.

To determine the functional distinctions between ACx ET and IT L5 neurons, we first performed two-photon calcium imaging from the apical dendrites of both types of neurons during a stationary-boundary auditory categorization task without rule-switching^38^. To image IT L5 neurons, we expressed GCaMP6s using Tlx3-Cre mice to specifically label these neurons. To image ET L5 neurons, we used a combinatorial viral expression strategy by injecting Retro-AAV-Cre in the intermedial and deep layer of SC, and injecting AAV-hSyn-Flex-Gcamp6s-WPRE-SV40 in the ACx in the same hemisphere (Extended Data Fig. 2; see Methods). We observed dendritic activity showing representations for sensory and category choice information in both ET and IT L5 neurons, but to different degrees. Linear regression and ANOVA showed that, over the population, the ET dendrites encode more choice information than the IT dendrites. Conversely, the IT dendrites encode more sensory information than the ET dendrites (Extended Data Fig. 2l and 2m). This distinction is corroborated by population decoding analysis using a support vector machine (SVM) with linear kernel across different time epochs of the behavioral trial, showing greater decoding accuracy for choices after sound stimulus delivery in ET dendrites than in IT dendrites (Extended Data Fig. 2p, q).

To further determine the causal roles of the L5 ET and IT neurons in the stationary-boundary auditory categorization task, we performed chemogenetic silencing on these two types of neurons respectively. We expressed hM4Di locally in the ACx with cell-type specificity in the two types of L5 neurons, respectively, in different groups of mice. To silence L5 ET or IT neurons in the ACx during task performance, we administered CNO (intraperitoneal injection) prior to each testing behavioral session (Extended Data Fig. 3; see Methods). As a control, we injected saline in alternating sessions in the same mice. We found that in mice with hM4Di expressed in ACx L5 IT neurons, the performance of auditory categorization was not significantly different between sessions with CNO injection and those with saline injection (Extended Data Fig. 3a-d). In contrast, in mice with hM4Di expressed in ACx L5 ET neurons, CNO injection significantly impaired the performance of auditory categorization (Extended Data Fig. 3e-h). This result suggests that L5 ET neurons play an essential role in our behavioral task. This is consistent with the anatomical and functional properties of L5 ET neurons, which extensively project to subcortical areas and show strong choice coding^45–48^. We thus focused our further investigation on how the dendritic activity of L5 ET neurons contributes to the computation of rule-dependent flexible decision-making in our RSFC task.

### Differential representations for categorization rule in the dendritic and somatic activity

To understand the sub-neuronal compartmentalization of task-relevant information during flexible decision-making, we optimized the imaging conditions to record calcium signals from the apical tuft dendrites, apical trunks, and the soma of L5 ET neurons in mice trained to perform the RSFC task (Fig. 1l,m; see Methods). We found that during this behavior, activity in the apical dendrites showed differential responses under the two task rules in both the reversing and non-reversing trials (Fig. 2a, b; Extended Data Fig. 4a-c). During the reversing trials, where the choice is different for the same set of reversing stimuli under the two rules, the response difference could reflect either rule difference or the choice difference. But during the non-reversing trials, where both the choice and stimuli were unchanged across the two rules, the response difference can be primarily attributed to the rule difference (Fig. 2c, d). Using the receiver operating characteristic (ROC) analysis, we quantified the selectivity for task rules in dendritic activity using non-reversing trials (see Methods). For the dendritic activity shown in Fig. 2b, we found a highly discriminative value of the area under ROC curve (auROC = 0.81, Fig. 2e), indicating a strong selectivity for task rules. We constructed a rule selectivity index (SI-rule) using the auROC value to represent the selectivity for either rule 1 (SI-rule > 0) or rule 2 (SI-rule < 0). For each ROI, the statistical significance of the SI-rule was determined by comparing to the distribution of auROC values from randomly shuffled data (see Methods). Overall, substantial proportions of dendritic ROIs showed significant selectivity to categorization rules (Fig. 2f; Extended Data Fig. 4d).

**Fig. 2.**
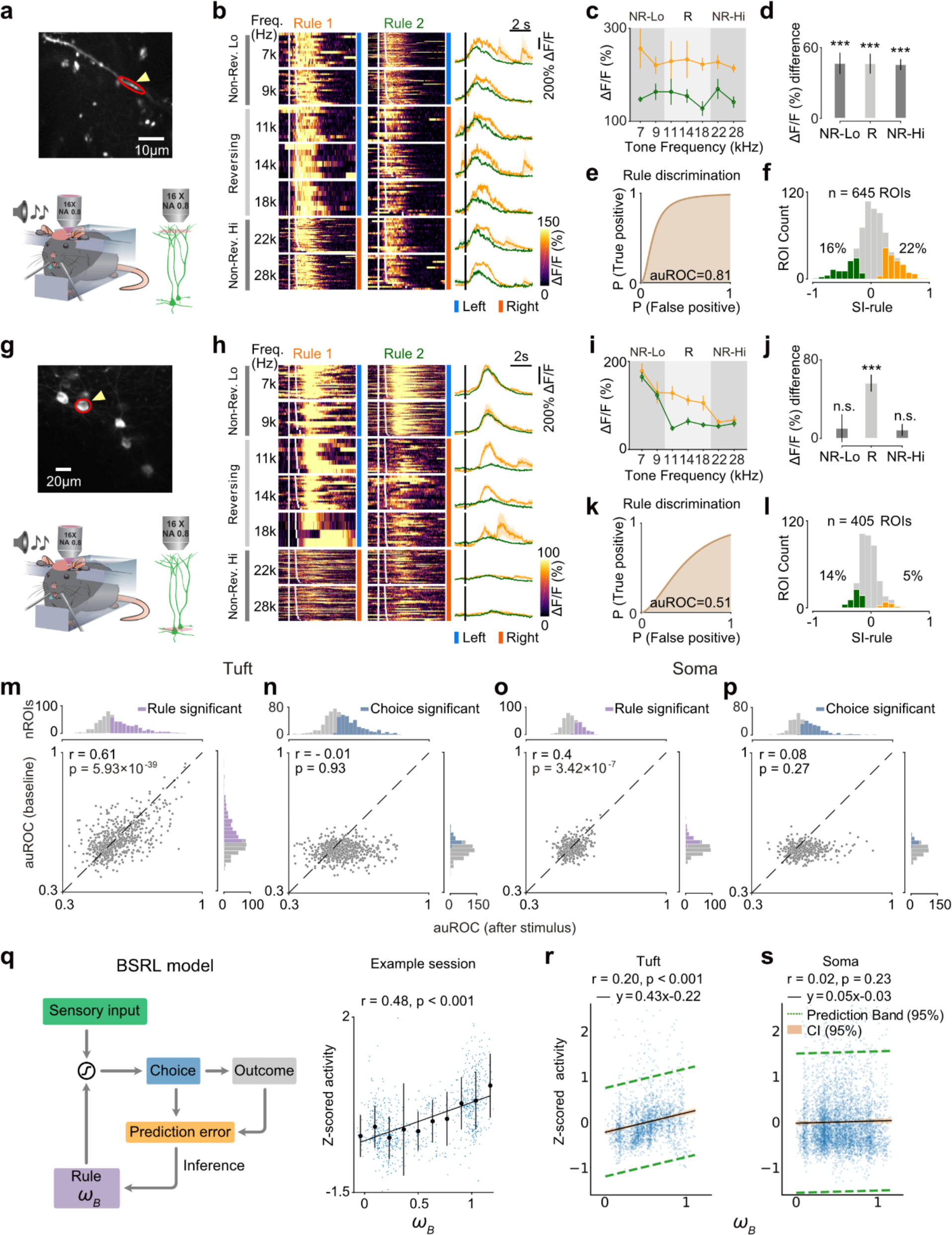
Categorization rule is preferentially encoded in apical dendrites of L5 ET neurons. **a**, Example two-photon image from the tuft dendrites of ACx L5 ET neurons acquired during task performance. Focal plane depth, ∼ 45 μm from pia. **b**, Color raster showing trial-by-trial Ca^2+^ signals from an apical tuft ROI indicated in **a**, in a typical behavioral session. Each row represents a single trial aligned to stimulus onset (first white line). Repeated trials of each tone frequency are grouped, and sorted according to trial types. Trials under the two rules are separated into the left and right columns. Trials within each frequency group are sorted by the answer lick time. Reversing and non-reversing trials are indicated by the light and dark gray vertical bars on the left respectively. Traces on the right show the averaged Ca^2+^ signals across trials within each stimulus condition under the two rules. Shades represent the SEM. **c**, Average responses from **b** plotted against tone frequencies under the two rules (see Methods). Error bars represent SEM. **d**, Activity difference between rule 1 and rule 2 for reversing and non-reversing trials in **b**, Wilcoxon rank-sum test, ***, p < 0.001. **e**, The ROC curve for the responses in **b** using correct non-reversing trials. The area under the ROC curve (auROC) indicates the degree of rule discrimination. **f**, Distribution of rule selectivity index (SI-rule) computed using ROC analysis (see Methods) from all tuft ROIs. Positive and negative values correspond to selectivity for rule 1 and rule 2, respectively. Yellow and green colors indicate the proportions with significant selectivity (p < 0.05, permutation test, n = 1000 random permutations). Gray, not significant. **g**, Example two-photon image of ET somas. **h**, Ca^2+^ signals as in **b**, from an example soma ROI indicated in **g**. **i**, Average responses from **h** as in **c**. **j**, Activity difference between rule 1 and rule 2 for data in **h**. **k**, The auROC for the responses in **h** for rule discrimination. **l**, Distribution of SI-rule in soma as in **f**. **m**, Correlation of auROC values for rule discrimination before and after stimulus onset in all dendritic tuft ROIs. Each dot represents auROC values from one ROI. X-axis, auROC value computed using responses after stimulus onset; y-axis, auROC value computed using calcium signals before stimulus onset (see Methods). Histograms on the top and right represent the distribution of auROC values with color indicating ROIs with significant rule selectivity (p < 0.05, permutation test, n = 1000 random permutations). **n**, Correlation of auROC values for choice discrimination before and after stimulus similar as in **m**. **o**, Similar as in **m** for all somatic ROIs. **p**, Similar as in **n** for somatic ROIs. **q**, Left, Schematic showing the BSRL model for calculation of the rule estimate. Right, Correlation between activity from rule selective tuft ROIs in an example imaging field and the task rule estimate from the BSRL model. Individual dots (blue) represent data from single trials. Correlation between trial-by-trial activity from example session rule selective tuft ROIs and rule estimate (*r* = 0.48, p < 0.001, 437 trials from 64 dendritic ROIs). Black solid circles with error bars are binned average for visualization, mean ± SEM. **r**, Correlation between trial-by-trial activity from all rule selective tuft ROIs and rule estimate (*r* = 0.20, *slope* = 0.43, p < 0.001, 2353 trials from 249 dendritic ROIs). **s**, Correlation between trial-by-trial activity from rule selective somatic ROIs and the task rule estimate (*r* = 0.02, *slope* = 0.05, p = 0.23, 5175 trials from 78 ROIs).

We next examined the task rule representation in the somatic activity of L5 ET neurons. The somatic responses of an example neuron shown in Fig. 2g, h exhibited prominent selectivity for category choice, as indicated by the pronounced response differences between different choices in reversing trials upon categorization rule change. However, in this example neuron, the responses in non-reversing trials are similar across different categorization rules (Fig. 2i, j), suggesting a lack of clear relationship with task rules. ROC analysis using non-reversing trials also yielded an insignificant discriminative value (auROC = 0.51, Fig. 2k). Overall, a smaller proportion of somatic ROIs showed significant rule selectivity (5% for rule 1 and 14% for rule 2, 405 neurons from 5 mice; Fig. 2l; Extended Data Fig. 5a). Within the rule-selective dendritic and somatic ROIs, the rule-related response differences in non-reversing trials are significantly less in soma comparing to dendrites (Extended Data Fig. 5b). The discriminability as measured by auROC is significantly higher in dendritic activity (auROC = 0.70 ± 0.0054), than in somatic activity (auROC = 0.63 ± 0.0047, *p* < 0.001, Wilcoxon rank-sum test; Extended Data Fig. 5c).

The rule inference during the task should be an ongoing process that persists beyond the sensory stimulus period in each trial. While the most prominent difference in dendritic activity between different task rules appeared to be after stimulus delivery (Fig. 2b), this could be partly due to the rule-related modulation riding on top of or being amplified by sensory evoked responses. We wondered whether the rule information could still be detected during the baseline period before stimulus presentation in each trial. We therefore performed ROC analysis on the rule coding using both the pre-stimulus (baseline) activity and post-stimulus activity. We found that for dendritic activity, a substantial proportion of ROIs showed significant rule coding both during baseline and post-stimulus periods, exhibiting a strong correlation across the population (Fig. 2m, r = 0.61, see also Extended Data Fig. 4e). Such persistent rule coding across baseline and post-stimulus periods is to a lesser extent in somatic activity (Fig. 2o, r = 0.4, p < 0.001 compared to the tuft ROIs, Fisher’s Z-transformation). In contrast to the persistence of rule information, choice information could only be decoded after stimulus presentation but not in the baseline phase both in dendritic and somatic ROIs (Fig. 2n, p; Extended Data Fig. 4f). In addition, choice information appears to be represented globally by both dendritic and somatic activity, for which the auROC values were not significantly different between dendritic and somatic compartments (Extended Data Fig. 5d).

It should be noted that the categorization rule is a hidden task variable that the subject must estimate based on the behavioral outcomes on a trial-by-trial basis^36^. The rule-selective neuronal activity could encode the subjective estimation of task rule. To establish the relationship between dendritic activity and the subjective rule inference, we used the belief-state reinforcement learning (BSRL) model to fit the behavioral data and computed the trial-by-trial rule estimate as an approximation of subjective rule inference (see Methods)^36^. We then examined the trial-by-trial correlation between the rule estimate and the calcium signals in the dendritic or somatic ROIs that showed significant rule selectivity (based on auROC values). This analysis revealed a significant correlation between dendritic activity and the rule estimate (Fig. 2q, r; *r* = 0.20, *p* < 0.001; see also Extended Data Fig. 4g). By contrast, in the somatic ROIs that showed significant auROC values, we did not find a significant trial-by-trial correlation between responses and rule estimate (*r* = 0.02, p = 0.23, Fig. 2s). Together, these results suggest that the representation of the subjectively inferred task rule is largely compartmentalized in the dendritic domains.

### Dendritic compartmentalization of multiple task variables

Our finding that the distal dendritic tuft receives and compartmentalizes task rule information is consistent with established knowledge about cortical circuits, including lamina-specific information flow and sub-neuronal input organization^3,11,13,34^. To systematically examine the sub-neuronal compartmental organization of behaviorally relevant information in our flexible categorization task, we used a generalized linear model (GLM) to predict the dendritic and somatic activity with the task and cognitive variables as predictors^24,49^. These predictors include sensory, choice, rule estimate, and prediction error (Fig. 3a, see Methods). The sensory variables are the different tone frequencies, while the choice variables are the categorical left and right choices measured from each trial. The rule estimate and prediction error are continuous variables calculated from the BSRL model. This encoding model captured a substantial proportion of the variance in both dendritic and somatic activities (Extended Data Fig. 6c-e; see Methods).

**Fig. 3.**
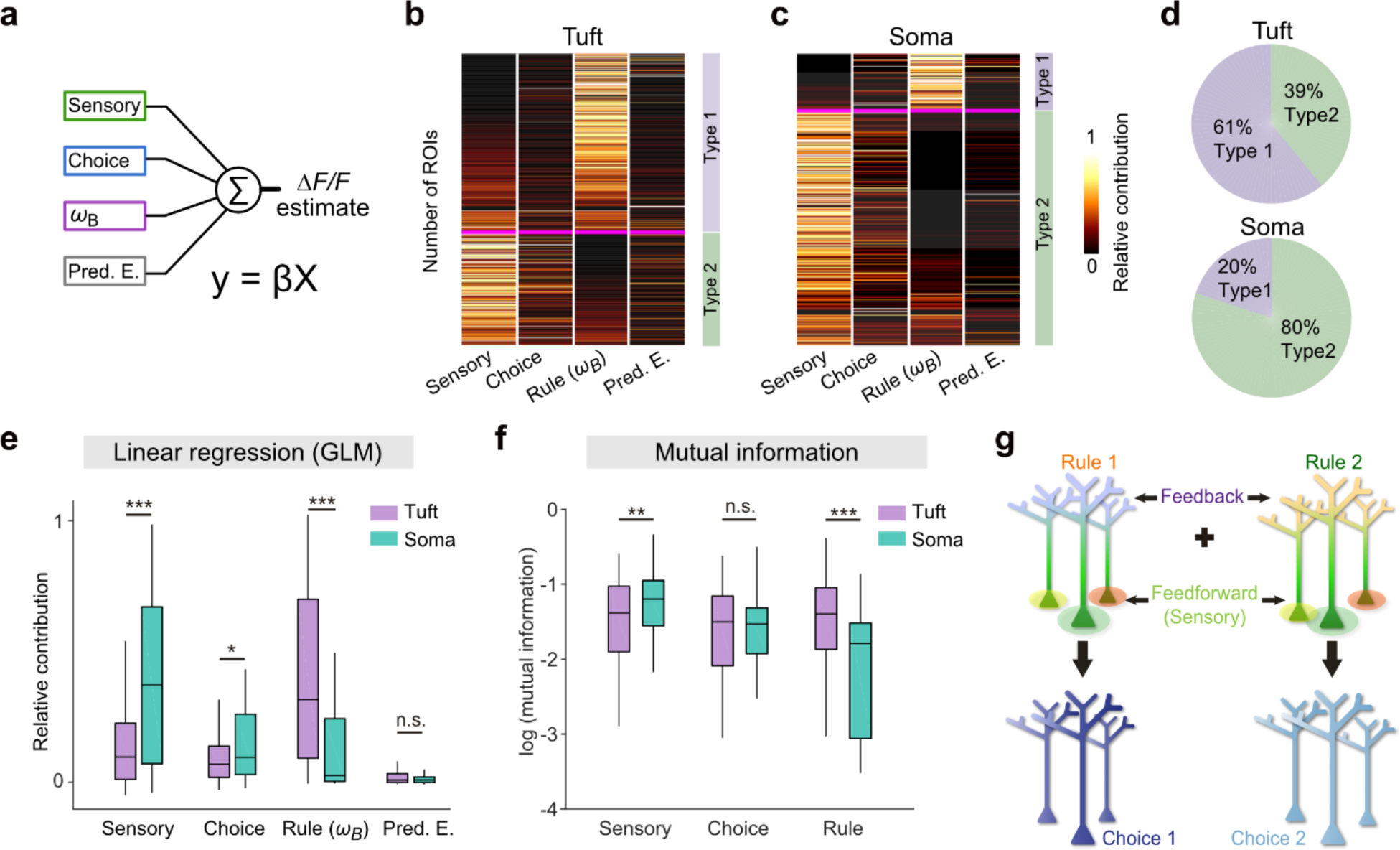
Distinct encoding of task variables by dendritic and somatic activity. **a**, Modeling the contribution of the behavioral variables to Ca^2+^ activity using a generalized linear model (GLM). The encoding of different task variables by the dendritic or somatic activity was determined based on the contribution of each variable (sensory, choice, categorization rule estimate and prediction error, Pred. E.) to the dendritic or somatic activity, quantified using the relative contribution to the explained variance of the calcium signals (see Methods). **b** and **c**, Clustering of all dendritic **b** and somatic **c** ROIs based on the relative contribution of different variables using the hierarchical-linkage-cosine method. Two main types (Type 1 and Type 2) were obtained for both dendritic and somatic ROIs (see Methods). **d**, Distinct composition of Type 1 and Type 2 in dendritic and somatic ROIs. Type 1: 395/645 in dendrites, 80/405 in somata; Type 2, 250/645 in dendrites, 325/405 in somata. **e**, Summary of relative contributions to dendritic and somatic ROIs by different variables. ROIs with a total explained variance > 0.1 were included. Comparison was made between dendritic ROIs (n = 174) and somatic ROIs (n = 93) for sensory (p = 7.49 × 10^-9^), choice (p = 1.18 × 10^-2^), rule (p = 1.09 × 10^-8^), and prediction error (p = 0.53), Wilcoxon rank-sum test, ***, p < 0.001; *, p < 0.05; n.s., not significant. **f**, Summary of mutual information (MI) between activity and different variables in dendritic tuft (n = 174) and somas (n = 93) as in **e**. MIs are compared between tuft and soma for sensory (p = 0.009), choice (p = 0.73), and rule (p = 1.87 × 10^-7^), Wilcoxon rank-sum test, ***, p < 0.001; **, p < 0.01; n.s., not significant. **g**, Schematic showing the hypothetical model where subcellular compartmentalized representations could be transformed into rule-dependent choice output via dendritic integrative mechanisms.

To characterize the overall difference in how dendrites and somas encode various task variables, we performed a clustering analysis on dendritic and somatic ROIs based on the variance explained by different task variables (see Methods). We found that both the dendritic and somatic ROIs could be classified into two primary distinct types, each displaying almost opposite preferences in the encoding for sensory stimuli and subjective rule estimate (Fig. 3b, c; Extended Data Fig. 6a). The first type (Type 1) of ROIs strongly encodes task rule information while weakly encodes sensory information. Conversely, the second type (Type 2) strongly encodes sensory information but weakly encodes task rule information. Notably, the composition of the Type 1 and Type 2 ROIs exhibits remarkable differences between dendrites and somas. Type 1 (rule dominant) accounts for 61% of the imaged dendritic tuft ROIs, 42% of the trunk ROIs, and only 20% of the somatic ROIs. In contrast, Type 2 (sensory dominant) constitutes 80% of the somatic ROIs, 58% of the trunk ROIs, and 39% of tuft ROIs (Fig. 3d; Extended Data Fig. 6b). Such representational differences imply a clear trend of directional information flow across sub-neuronal compartments: task rule information is mostly represented in dendritic tuft, while exhibiting a substantial attenuation when spreading to the soma, whereas sensory information is mostly represented in the soma, while showing significant reduction when propagating towards the distal dendrites.

We further compared the encoding for different variables between the dendritic and somatic ROIs based on their relative contributions to the explained variance in the GLM. Across the population, somatic ROIs encode significantly more sensory information than dendritic ROIs, whereas dendritic ROIs encode significantly more rule information than somatic ROIs (Fig. 3e; Extended Data Fig. 6f-i). Choice information, however, tends to be globally represented, with a slightly higher degree in somas than in dendrites (Fig. 3e).

In addition to GLM-based regression analysis, we also estimated the mutual information (MI)^50–52^ between calcium activity and various task variables (see Methods). Consistent with the trends observed in the GLM analysis, dendritic activity shows a higher level of estimated MI for task rule (MI-rule) compared to somatic activity, while somatic activity exhibits a higher level of estimated MI for sensory stimuli (MI-sensory) than dendritic activity (Fig. 3f; Extended Data Fig. 6j). The MI for choice (MI-choice) is found at the similar level in dendritic and somatic ROIs (Fig. 3f; Extended Data Fig. 6j).

Together, these results demonstrate that cortical L5 ET neurons exhibit sub-neuronal compartmentalization in coding for key sensory and cognitive variables, with distinct representations between dendrites and somas during a cognitive task. This sub-neuronal compartmentalization of behavioral and cognitive variables may enable dendritic nonlinear integration^7,20^ to implement a rule-dependent flexible stimulus categorization (Fig. 3g).

### Single-neuron measurements of compartmental representations

We next asked whether the dendritic rule information is conveyed via local dendritic inputs or inherited from somatic backpropagating action potentials (bAPs, Fig. 4a). We reasoned that if there was localized dendritic activity within individual branches encoding rule information, it would support the possibility of direct dendritic inputs carrying rule information. Indeed, we found that in some cases, highly localized activity in subregions of dendritic branches exhibited significant rule coding (Fig. 4b-d), but the widespread activity across multiple branches often did not (Fig. 4e). This isolated dendritic activity appears to arise from direct dendritic inputs that have not been integrated with backpropagating somatic activities, suggesting that rule information could be conveyed by direct inputs to the distal dendrites.

**Fig. 4.**
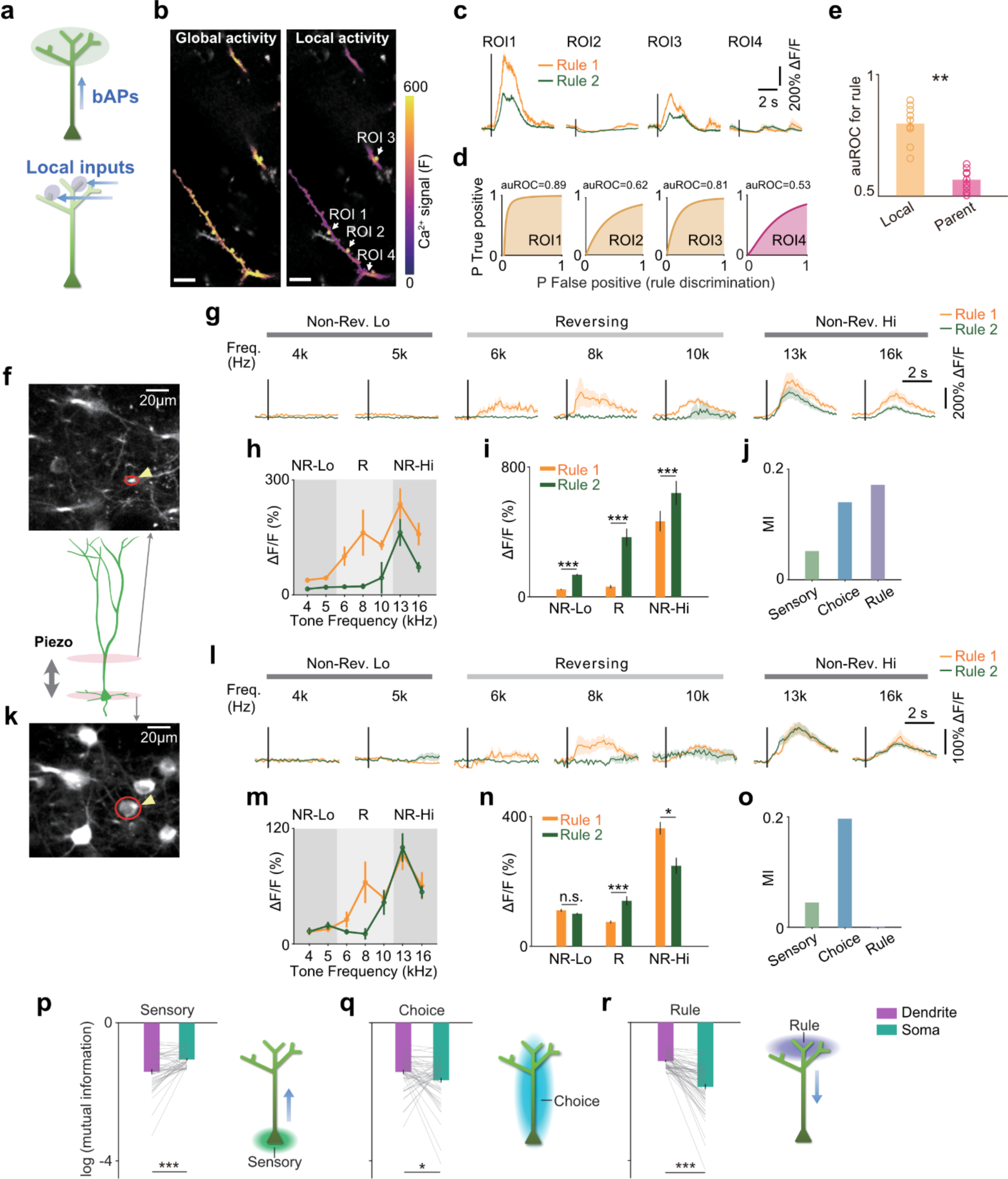
Compartmentalized representations within single neurons. **a**, Schematic showing two alternative hypotheses with dendritic rule information inherited from bAPs or arising from local inputs. **b**, Example local and global dendritic activity. Left, color-coded Ca^2+^ signals in an example trial showing correlated activity across multiple tuft branches. Right, color-coded Ca^2+^ signals in an example trial showing localized activity. Color represents the maximum pixel values during the trial. Scale bars, 10 μm. (see Extended Data Movie S1-S2). **c**, Ca^2+^ signals from multiple trials from the local dendritic spots or nearby branch segments as indicated by arrowheads in **b**. **d**, auROC for rule discriminability of corresponding ROIs. **e**, Summary of auROC values compared between rule selective local sub-branch dendritic activity and the activity from the parent branches across visually identified neurons (9 out of 23 visually identified neurons with sub-branch activity exhibited local rule coding, Wilcoxon sign-rank test, p = 0.0039. **f**, Example of semi-simultaneous two-plane imaging of the same L5 ET neurons during task performance from a dendritic region. **g**, Traces show the averaged Ca^2+^ signals across trials within each stimulus under the two rules from dendritic ROI shown in **f** with the red circle. Shades represent the SEM. **h**, Average responses from **g** plotted against tone frequencies under the two rules. Error bars represent SEM. **i**, Ca^2+^ signals in the example dendritic ROI in **g**, compared between rule 1 and rule 2 for reversing (R) and non-reversing trials (low frequency, NR-Lo; high frequency, NR-Hi). Wilcoxon rank-sum test, ***, p < 0.001. **j,** Mutual information for different variables from dendritic activity in **g**. **k**, Example field from the somatic region simultaneously acquired with same neuron dendritic region shown in **f**. The red circle labels a somatic ROI from the same neuron with the identified dendrite as indicated in **f**. **l**, Traces show the averaged Ca^2+^ signals across trials within each stimulus under the two rules from somatic ROI shown in **k** with the red circle. **m**, Average responses from **l** plotted against tone frequencies under the two rules. Error bars represent SEM. **n**, Ca^2+^ signals in the example somatic ROI in **k**, compared between rule 1 and rule 2 for reversing (R) and non-reversing trials (low frequency, NR-Lo; high frequency, NR-Hi). Wilcoxon rank-sum test, ***, p < 0.001; *, p < 0.05; n.s., not significant. **o**, Mutual information as in **j** for the somatic ROI. **p**, Summarized mutual information (MI > 0) for sensory stimuli (MI-sensory) compared between all somas and dendrites within the same individual neurons (Wilcoxon sign-rank test, p = 1.13 × 10^-5^, n = 47 neurons with significant somatic MI-sensory). **q**, Mutual information (MI > 0) for choices (MI-choice) compared between all somas and dendrites within the same neurons (Wilcoxon sign-rank test, p = 0.04, n = 52 neurons with significant MI-choice). **r**, Mutual information (MI > 0) for rules (MI-rule) compared between all somas and dendrites within all same neurons (Wilcoxon sign-rank test, p = 1.43 × 10^-11^, n = 62 neurons with significant dendritic MI-rule). Error bars represent SEM. *, p < 0.05, **, p < 0.01, ***, p < 0.001.

We next sought to examine the dendritic and somatic compartmentalization of task information within the same neurons. We performed semi-simultaneous two-plane calcium imaging from both somatic and dendritic regions (including tuft and trunk) during task performance. We then identified dendritic and somatic ROIs belonging to the same neurons (Fig. 4f, k; see Methods), and carried out pair-wise comparison of the information coding between the dendritic and somatic ROIs within the same neurons. As illustrated in the example dendritic and somatic ROIs from a single neuron, the dendritic activity in the non-reversing trials is modulated by rules and exhibits significant mutual information (MI) for choice and rule (Fig. 4g-j), while the somatic activity primarily shows choice-related modulation in the reversing trials and significant choice MI (Fig. 4l-o).

To examine compartmentalization of sensory information within single neurons, we identified neurons with significant sensory MI in somatic activity (p < 0.05, permutation test), and compared sensory MI between soma and dendrites (Fig. 4p and Extended Data Fig. 7). We found that sensory MI was significantly higher in the somatic activity than in the dendritic activity (Fig. 4p, p = 1.13×10^-5^, n = 47), indicating somatic compartmentalization of sensory information. For the dendritic compartmentalization of rule information, we identified neurons with dendritic ROIs showing significant rule MI (p < 0.05, permutation test), and compared rule MI in dendritic and somatic activity in the same neurons. The rule MI in the dendritic activity was significantly higher than in somatic activity (Fig. 4r, p = 1.43×10^-11^, n = 62), supporting the dendritic compartmentalization of rule information. For choice information, we examined neurons with significant choice MI in either dendrites or somas and found a slightly higher level of choice MI in the somatic activity (Fig. 4q, p = 0.04, n = 52). These results indicate that, consistent with our population data analysis, within single L5 ET neurons, the apical dendrites primarily compartmentalize rule information, whereas the soma mainly compartmentalizes sensory information.

### Contributions of dendritic activity to the computation for rule-based categorization

Based on the dendritic coincidence detection model^7,9,20,34^, our results are consistent with the hypothesis that the integration of dendritic rule information with the backpropagated sensory information could result in a rule-based stimulus categorization. This hypothesis predicts that the dendritic activity is necessary for somatic coding for rule-specific categorical choices. To test this prediction, we simultaneously inhibited the dendritic activity while imaging the dendritic or somatic activity of L5 ET neurons. To inhibit dendritic activity, we leveraged the property of inhibitory somatostatin-expressing (SST) interneurons which preferentially target distal apical dendrites^53–57^. We expressed a red-light sensitive channel rhodopsin (ChrimsonR) in SST neurons using SST-Cre mice. To activate SST neurons in the same region of two-photon imaging, the red laser light (635 nm) was delivered through the objective via a custom-designed light path incorporated into the 2P microscope (Fig. 5a; Methods). To examine the effect of optogenetic stimulation within the same session, the optogenetic photostimulation was delivered in a subset (∼50%) of trials interleaved with control trials (without photostimulation). To reduce the potential optical influence on imaging by the optogenetic light illumination, the red laser (635 nm) was delivered only during the fly-back period of each line scan such that photostimulation was always outside of the acquired 2P images (Fig. 5b; Methods). Photostimulation of SST neurons effectively reduced the dendritic coding for the task rule information, as evidenced by the significantly decreased auROC values (Fig. 5c) and the reduced relative contributions of rule information to the explained variance quantified using GLM (Extended Data Fig. 9).

**Fig. 5.**
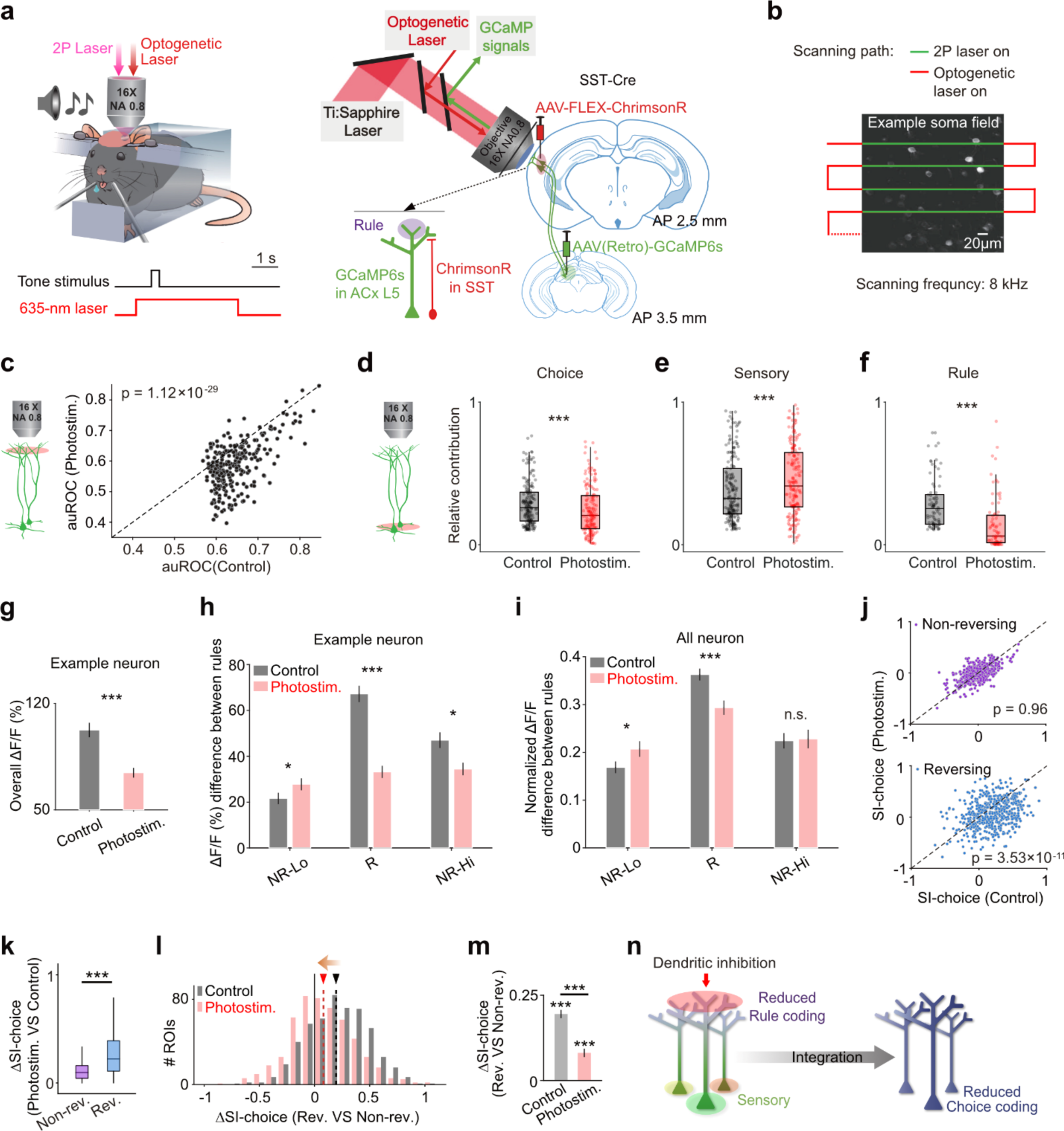
Dendritic inhibition disrupts somatic choice output. **a**, Schematic showing the experimental configuration of simultaneous photostimulation of SST neurons and imaging of L5 ET neurons in the same imaging field in ACx. ChrimsonR was expressed in ACx SST neurons and GCaMP6s was expressed in SC projecting ACx neurons using the retrograde labeling method (Methods). On a photostimulation trial, the red-light illumination began 0.5 s before sound stimulus onset and lasted for 4 s. **b**, To avoid artifacts from optogenetic illumination during imaging, the optogenetic laser was delivered outside of image acquisition (see Methods). **c**, Discrimination of rule (auROC) by individual dendritic ROIs compared between control trials and photostimulation trials (Wilcoxon signed-rank test, p < 0.001, n = 262 dendritic ROIs showing significant rule selectivity under the control condition). **d**-**f**, Somatic coding for choices **d**, sensory stimuli **e**, and rules **f**, measured as the relative contribution to the explained variance in the GLM model, compared between photostimulation and control trials within individual neurons (choice selective neurons, p = 8.3 × 10^-5^, n = 172; sensory selective neurons, p = 5.81 × 10^-4^, n = 182; rule selective neurons, p = 1.17 × 10^-7^, n = 78, Wilcoxon signed-rank test; see Methods). **g**, Comparison of overall Calcium activity between control (295 trials) and photostimulation (268 trials) from an example soma of an example session. Wilcoxon rank-sum test, p = 1.35 × 10^-5^, ***, p < 0.001. **h**, Comparison of Calcium activity difference between control and photostimulation for example soma in **g**, Wilcoxon rank-sum test, ***, p < 0.001; *, p < 0.05. **i**, Comparison of activity difference under different rules between control and photostimulation for all soma ROIs normalized by each neuron’s max activity, n = 521, NR-Lo, p = 0.02; R, p = 6.36 × 10^-4^; NR-Hi, p = 0.81, paired t-test, ***, p < 0.001; *, p < 0.05; n.s., not significant. **j**, Choice selectivity index of somatic activity compared between control trials and photostimulation trials for non-reversing stimuli (upper, p = 0.96, n = 521, Wilcoxon signed-rank test) and reversing stimuli (lower, p = 3.53 × 10^-11^, n = 521, Wilcoxon signed-rank test). **k**, Changes in choice selectivity index (ΔSI-choice) following photostimulation, compared between reversing and non-reversing trials (p = 2 × 10^-46^, n = 521, Wilcoxon signed-rank test). **l**, Distributions of the difference in somatic choice selectivity between reversing trials and non-reversing trials, under photostimulation (red) and control (gray) conditions. Dashed lines indicate the mean value. Photostimulation shifted the distribution towards zero as indicated by the orange arrow. **m**, Summary of the data shown in **l** between photostimulation and control conditions. Wilcoxon signed-rank test, p = 8.73 × 10^-16^, n = 521. **n**, Schematic summarization of the effect of dendritic inhibition on dendritic rule coding and somatic choice coding.

To assess the impact of dendritic inhibition on somatic output, we imaged L5 ET somas during photostimulation. For neurons whose somatic activity could be significantly explained by the regression model, we identified choice selective, sensory selective, and rule selective neurons, and compared the relative contributions to the explained variance by corresponding variables between control and photostimulation trials (Methods). We found that somatic coding for category choices was significantly reduced following dendritic inhibition (Fig. 5d), suggesting that dendritic activity was required for the somatic output of the stimulus categorization. Somatic coding for rule information was also significantly reduced following dendritic inhibition (Fig. 5f), consistent with the notion that somatic rule information mainly originates from the dendrites. By contrast, the relative contribution from sensory stimuli was slightly increased following dendritic inhibition (Fig. 5e), suggesting that somatic sensory responses could be less susceptible to dendritic inhibition. The slight increase could result from a relatively greater reduction in the choice and rule information. Together, these results support that dendritic activity contributes to somatic coding for choices.

To further verify the contribution of dendritic activity to the computation of rule-based stimulus categorization, we separately examined the dendritic inhibition effects on reversing and non-reversing trials. We reasoned that due to distinct stimulus-action-reward contingencies in reversing and non-reversing trials over the behavioral training, the strength of stimulus-action associations can grow differently for the two types of stimuli. For non-reversing stimuli, the fixed sensory to motor mappings could be substantially strengthened over training such that strong choice-related output could be directly elicited by the sensory input without requiring the integration of dendritic rule information. For reversing stimuli, however, the sensory to motor mappings constantly change, contingent on the categorization rules, such that only through nonlinear integration with dendritic rule information, could the sensory input be transformed to strong choice output. Indeed, following dendritic inhibition, the somatic choice selectivity showed greater changes in reversing trials than in non-reversing trials (Fig. 5g-k). In addition, we observed that somatic activity typically exhibited a greater degree of choice coding in reversing trials compared to non-reversing trials, presumably reflecting dendritic task rule modulation during reversing trials (Fig. 5l). This difference was significantly reduced after dendritic inhibition by photostimulation, with the mean difference approaching zero (Fig. 5l, m). Together, these results indicate that reducing dendritic rule coding via optogenetic inhibition disrupted rule-dependent somatic choice coding (Fig. 5n).

## Discussion

Dendritic computation has long been proposed to be the central mechanism for neuronal information processing^4,6,7,13,29,34,35,58,59^. However, direct evidence supporting dendritic computation in complex behaviors involving cognitive level processing has been lacking. On one hand, although dendritic activity has been studied in certain sensorimotor behaviors, the prevailing observations were that the soma and dendrites are strongly coupled, showing primarily global dendritic spikes or bursting somatic activity^19–22,60^. On the other hand, dendritic specific activity was often observed with passive sensory stimulation^15–17,61^, or during certain motor behaviors^23,24,62^ without further understanding its role in cognitive-level computations.

How dendrites compartmentalize and integrate task-related information to implement computations underlying cognitive functions remains largely unknown. To tackle this problem, we created a task condition where we could isolate the cognitive variable of task rule inference, by which animals solve a complex rule-switching flexible categorization task^36^. Using two-photon imaging to record activity in sub-neuronal dendritic and somatic compartments under this task condition, we discovered, to our knowledge for the first time, the dendritic and somatic compartmentalization of both internally generated cognitive information and task-specific sensory and motor variables. Moreover, we demonstrate that the compartmentalized dendritic representations play a crucial role in the somatic output of rule-based flexible categorization. Together with the prior notions of dendritic nonlinear integration^7,9,10,20,34^, our findings suggest a dendritic computational schema that the apical dendrites of L5 pyramidal neurons compartmentalize internal cognitive information of task rules, which nonlinearly integrates with somatic sensory inputs to generate flexible choice output. This dendritic computation is task relevant, as it is selectively involved in rule-dependent dynamic remapping of choices for reversing stimuli, but is not required for non-reversing stimuli with fixed choice associations. A plausible mechanism is that, following task training, direct sensory inputs from non-reversing stimuli become sufficiently strong to directly drive bursting activity to influence choices, whereas reversing stimuli require rule-dependent nonlinear dendritic integration to produce flexible choice output.

Our results suggest that the internal cognitive information of rule inference was carried by inputs to the distal dendrites (Fig. 4). The spread of these dendritic signals towards somatic regions could be subject to dendritic filtering and local inhibition, such that these signals per se could hardly generate somatic action potentials. Consequently, the task rule information could be reliably decoded from dendritic GCaMP6s Ca^2+^ signals, but is largely undetectable in somatic GCaMP6s Ca^2+^ signals which mainly reflect somatic APs (Fig. 3d; Extended Data Fig. 6). On the other hand, sensory inputs to the proximal somatic regions can generate somatic APs, which could backpropagate to the apical dendrites, and account for the detectable but less prevalent sensory information in the dendrites with a dependence on distances from somas (Fig. 3d; Extended Data Fig. 6b).

The dendritic coincidence detection model suggests that dendritic activity is essential in generating salient somatic output. Here we tested this by manipulating dendritic activity while measuring the somatic output of the categorization computation. We exploited the sub-neuronal target specificity of the SST+ interneurons^32,53,54,56,57^, and combined a custom-developed all optical interrogation system to address this question. The SST+ interneurons, especially the deep layer SST+ interneurons known as the Martinotti interneurons preferentially target distal dendrites of pyramidal neurons^53,54,57^. Activation of these SST+ interneurons led to a significant reduction in rule information coding in the dendrites (Fig. 5 and Extended Data Fig. 9), supporting the effectiveness of this method in manipulating dendritic activity. Due to further diversity within SST+ neurons, our manipulation does not rule out the possibility of influencing other postsynaptic targets of SST+ neurons in the local circuits. Nevertheless, we took additional caution by targeting viral expression in deep layer SST+ and choosing imaging/manipulation region largely devoid of SST+ soma in the upper layer. Future development of new tools such as those targeting dendritic ion channels would be important to further improve the effectiveness and specificity of dendritic activity manipulation.

It has been increasingly recognized that distinct types of cortical L5 pyramidal neurons are associated with distinct functions^24,48,63^. However, a projectome-based systematic characterization of L5 neurons has not yet been applied to investigate dendritic function during behavior. In this study, we utilized whole-brain projectome analysis based on complete single-neuron morphological reconstruction, which we developed previously^43^, to investigate two main types of L5 neurons in the ACx: SC projecting ET neurons and the contralateral projecting IT neurons in (Extended Data Fig. 1a-f). The two types of neurons demonstrated distinct brain-wide projection patterns and dendritic morphologies (Extended Data Fig. 1g-h). Moreover, their dendrites exhibited distinct information coding properties, with IT dendrites primarily coding for sensory information and ET dendrites showing comparatively more category choice information (Extended Data Fig. 2). The preferential coding for sensory information in IT neurons, along with their contralateral cortical projection, suggests that these neurons may be primarily involved in processing of sensory features and coordinating sensory input from diverse spatial and temporal sources. However, advanced sensory processing appears to be non-essential in our current behavioral task, which emphasizes the complexity in task structure rather than in sensory features. Indeed, chemogenetic silencing of ACx L5 IT neurons did not produce any significant effects on the pure tone based auditory categorization task (Extended Data Fig. 3a-d). In contrast, the L5 ET neurons possess more extensive and larger apical dendritic trees and project widely to various subcortical brain regions, broadcasting the cortical output of choice-relevant auditory categorization. These structural features suggest that L5 ET neurons may play an essential role in integrating sensory and task-relevant information to influence behavioral output. Consistent with this notion, chemogenetic silencing of L5 ET neurons significantly impaired auditory categorization behavior (Extended Data Fig. 3e-h). Based on these systematic structural and functional characterizations of the major cortical L5 cell types, we focused on L5 ET neurons to investigate the role of dendritic integration in cognitive processing during rule-based flexible categorization.

The dendritic and somatic compartmentalization of different types of task-relevant information can also be manifested in a trend of graded representations along the apical dendrite-soma axis. Our data show that the proportion of rule preferred ROIs (Type I) is 61% for tuft, 42% for trunk and 20% for soma. Conversely, the proportion of sensory preferred ROIs (Type II) is 39% for tuft, 58% for trunk and 80% for soma (Fig. 3b-d, Extended Data Fig. 6a, b). While our current study focuses on the compartmental representations of different types of task information, future research may be directed to the investigation of the sources of the different types of inputs. The sensory information could primarily carried by the inputs from the thalamocortical projections^64^, e.g., the medial geniculate body (MGB), or from the local layer 4 neurons known to be the main recipients of thalamocortical input in the ascending sensory pathways^3,65^. For the rule information, there could be multiple possible sources. One is the long-distance input from higher order association cortical areas, e.g., the orbitofrontal cortex^36,66,67^, the posterior parietal cortex^37,43,66^, the secondary motor cortex^43,68^, or the higher order thalamic areas. Another possibility is that after task learning, the L2/3 neurons in ACx could undergo reorganization to represent rule-related stimulus category information^36,38^. These local L2/3 neurons are known to impinge on the apical dendrites of L5 neurons^65,69,70^, providing potential inputs carrying information of categorization rule. These possibilities should be the subjects for future investigation.

Our study provides a new viable approach to delineate circuit and cellular mechanisms involved in cognitive function. An interesting direction is to apply this approach to investigate the role of cortical inhibitory circuits in cognitive functions. For instance, while we used SST activation as a tool to inhibit dendritic activity, SST neurons could play a critical role in shaping dendritic activity and local population responses during behavior; the vasoactive intestinal peptide positive (VIP+) interneurons have been shown to mediate a gain modulation of sensory cortical responses via disinhibitory circuits^71–74;^ the parvalbumin positive (PV+) interneurons receive cortical feedback input and control the somatic spiking output^57,68^. Future studies may be directed to investigate the interactions between different subtypes of local inhibitory interneurons and dendritic functions during flexible behaviors.

In sum, we have discovered a dendritic computational mechanism in the cortical pyramidal neurons for realizing rule-inference based flexible decision-making, where internally derived rule information encoded in the apical dendritic activity nonlinearly integrate with somatic sensory input to produce dynamic output, presumably in the form of plateau potentials or bursting activity to drive flexible behavioral choices. This represents a fundamental circuit and neuronal mechanism underlying biological intelligence, and may additionally provide inspirations for developing more efficient computations in artificial system.

## RESOURCE AVAILABILITY

Further information and requests for resources and reagents should be directed to N.L.X.

## Materials availability

This study did not generate new unique reagents.

## Data and code availability

The data generated in this study to reproduce all the results have been deposited in Figshare DOI: 10.6084/m9.figshare.24968865

## Acknowledgments

We thank Dr. Mu-ming Poo for discussion on manuscript; Dr. Chunyu. A. Duan for discussion on data analysis; Zimu Li for help in data analysis; Keqing F. Xu for drawing schematics; Zhaomei Ying and Chun Xie for lab administration.

## Author contributions

Y.Z. and N.L.X. conceived the project and designed the experiments. Y.Z. performed the experiments and data analysis. Y.L. contributed to behavioral task design and behavioral training, and data analysis. L.C. helped in behavioral task development imaging method development. Q.L. helped in behavior training and imaging data collection. L.Z. helped in designing behavioral apparatus. Y.H. performed analysis on the projectome data. Y.X. helped in imaging data analysis. R.C. helped in chemogenetic experiments. L.D. helped in behavioral training and animal preparation. J.P. helped in imaging system designing and behavioral apparatus designing. J.D. helped in behavioral training and data analysis. Z.K.Z. helped in imaging data collection and data analysis. J.S. contributed to manuscript writing. Y.Z. and N.L.X. wrote the manuscript.

## Methods

### Experimental subjects

The Animal Care and Use Committee of the CAS Center for Excellence in Brain Science and Intelligence Technology, Chinese Academy of Sciences, approved the experimental procedures. Male C57BL/6J (SLAC), Tlx3-Cre (MMRRC, no. 041158-UCD), and SST-Cre (Jackson Laboratory, no. 013044) mice, aged 7–10 weeks at the beginning of the experiment, were the primary source of data. For two-photon imaging during stationary boundary auditory decision task, 4 Tlx3-Cre mice were used for imaging intratelencephalic (IT) L5 neurons, and 3 wild-type mice were used for imaging L5 ET neurons in ACx. For two-photon imaging during the flexible decision-making task, 5 wild-type mice were used for imaging L5 ET neurons. For experiments testing causal effect on behavior, 4 Tlx3-Cre mice and 5 wildtype mice were used for chemogenetic inhibition of L5 IT and ET neurons respectively. For two-photon imaging with optogenetic dendritic inhibition, 3 SST-Cre mice were used. On average each animal was used for 2-4 sessions.

### Surgery and virus injections

During surgery, mice were anesthetized with isoflurane (1-2%), with body temperature monitored and maintained at ∼37°C with a heating pad. The surgery details are as previously described^37^. To image ACx cortico-cortical projection neurons, AAV-hSyn-Flex-Gcamp6s-WPRE-SV40 virus (titer: 2-3×10^12^ i.u.; Shanghai Taitool Bioscience Co. Ltd.) was slowly injected (at a speed of 1 nl/5s, 50 nl per site, 2-3 injection sites per animal) into the auditory cortex (−2.6 mm AP, 4.5 mm ML from bregma, 0.7 mm below the dura) in Tlx3-Cre mice. To image ACx L5 ET neurons, AAV(Retro)-hSyn-Cre-mCherry was injected in three sites in SC (−3.5 mm AP, 0.6 mm ML, 2.0 mm DV from bregma; −3.5 mm AP, 1.0 mm ML, 2.0 mm DV from bregma; −4.0 mm AP, 1.0 mm ML, 2.0 mm DV from bregma) of wild-type mice, and AAV-hSyn-Flex-Gcamp6s-WPRE-SV40 virus was injected in the auditory cortex in the same animals. To image L5 ET neurons during dendritic photoinhibition, AAV(Retro)-hSyn-GCaMP6s was injected in SC of SST-Cre mice, and AAV-Syn-FLEX-ChrimsonR-tdTomato (2-3×10^12^ i.u.; Shanghai Taitool Bioscience Co. Ltd., Addgene: #62723) was injected in the ACx at a 45-degree angle and a depth of 0.8 mm below the dura to target the deep layer SST neurons (50 nl per site, 2 sites per animal).

For chemogenetic manipulation of L5 IT neurons, AAV2-hSyn-DIO-hM4D(Gi)-mCherry (1×10^13^ i.u.; Shanghai Taitool Bioscience Co. Ltd.) was bilaterally injected in the ACx of Tlx3-Cre mice. For chemogenetic manipulation of L5 ET neurons CAV2-Cre in TBS +15% glycerol (1×10^13^ i.u.; Shanghai Taitool Bioscience Co. Ltd.) was injected bilaterally in SC of wild-type mice, followed by AAV2-hSyn-DIO-hM4D(Gi)-mCherry injection in bilateral ACx. For retrograde labeling of the somata of L5 ET and IT neruons, AAV-Syn-FLEX-EGFP (50 nl/site) (1×10^13^ i.u.) was injected in Tlx3-Cre mice, and CTB-647 was injected in the ipsilateral SC. Mice were sacrificed 4 weeks after virus injection, and brain slices were imaged using an Olympus VS120 Virtual Slide Fluorescence Microscope.

For imaging window implantation, a craniotomy (3 mm in diameter) was made over the left auditory cortex after virus injection, with the dura intact. The imaging window consisted of a single-layered circular glass made from coverslip (200-300 μm thick) glued to a glass ring (outside diameter, 3 mm; inside diameter, 2 mm; height, 0.3 mm, from Guluo Glass Co. Ltd.), and the glass ring was then attached to a larger metal shim (stainless steel, outside diameter, 4.5 mm; inside diameter, 2.2 mm; thickness, 0.3 mm). The metal rings and glass were glued together using ultraviolet-cured optical adhesive (Norland Optical Adhesive 81). The glass window was inserted to the craniotomy contacting the dura, and metal ring was sealed to the skull with dental cement (Jet Repair Acrylic, Lang Dental Manufacturing). A titanium head-post was then attached to the skull with dental cement for head fixation. Mice were allowed to recover for at least 7 days before water restriction.

### Behavioral apparatus

To perform the auditory-based behavior, experiments were conducted in custom-designed sound-attenuating boxes, as described previously^37,38^. Briefly, mice were head-fixed with custom-made holders, with their bodies placed in an acrylic tube (25 mm in diameter). Two metal lick spouts were positioned in front of the mouse’s mouth for lick detection and water delivery. Licks were detected by a capacitive-sensing circuit board that was connected to the ports. The behavior protocol was controlled by a custom-developed real-time control system (PX-Behavior System), which included a custom-designed tone-generating module (TGM) and a microcontroller (Arduino MEGA 2560, IDE 1.5.6 or 1.8.2) for stimulus delivery and measurement of behavioral events. To deliver sound stimuli, the TGM sent specified waveforms to an amplifier (ZB1PS, Tucker-Davis Technologies) to drive a speaker (ES1, Tucker-Davis Technologies) placed on the right side of the mice. The sound system was calibrated with a free-field microphone (Type 4939, Brüel and Kjær) to ensure a loudness of approximately 70 dB SPL across all tested frequencies at the position of the mouse’s ear. 5 ms cosine ramps were applied to the rise and fall of all tones. Behavioral data were logged with custom-written software in Python (v2.7).

### Behavioral tasks

Mice were housed in groups of less than six per cage and subjected to a 12-hour reverse light/dark cycle (dark from 10:00 to 22:00) with all experimental procedures conducted during the dark phase. Mice had no history of previous experiments. Before training, water restriction and handling procedures outlined by Guo et al.^75^ were performed. Each mouse received 1 mL water per day and the body weight was monitored. After ∼7 days of water restriction, behavior training was started. Mice were allowed to perform the task until sated. During training days, each mouse received water during task performance, with supplements reaching approximately 1 mL/day. On non-training days, 1 mL of water was directly provided. Their body weight was measured and maintained at no less than 80% of the weight before water restriction.

The behavioral task is based on the auditory-guided two-alternative-forced-choice (2AFC) task^37,38,75^. Mice were first trained to discriminate two pure tone stimuli (duration, 300 ms) that were 2 octaves apart (7 kHz and 28 kHz, or 4 kHz and 16 kHz). The sound intensity was 70 or 75dB SPL for all frequencies. The initiation of each trial is not explicitly cued, and the animal needs to wait for the tone stimulus to occur. Mice need to respond within a 3 s answer following the stimulus offset by licking left or right lick port placed in front of the animal. Correct answers were defined as licking the left lick port in response to the lower frequency tone, or licking the right lick port in response to the higher frequency tone. Correct responses lead to the water valve open to dispense a small amount of water reward (∼6ul). A 2-4 s inter-trial interval (ITI) was imposed between trials. Error responses lead to a 2∼6 s in addition to ITI as a time-out punishment during which licking to the wrong side would reinitiate the time-out period. If mice made no response lick within the 3 s response window, the trial was defined as a ‘miss’ trial. Once mice reached a criterion of > 85% correct rate, the second training stage was started for testing categorization performance.

For the categorization task with stationary boundary (non-rule-switching, Extended Data Fig. 2 and 3), the tone stimulus in each trial was randomly chosen from 6 different frequencies (separated by linearly spaced intervals in octave, 7000 Hz, 9237 Hz, 12188 Hz, 16082 Hz, 21220 Hz, 28000 Hz). Reward contingency was based on the mid frequency (14 kHz) as the boundary, with correct answers defined as licking leftward when tone frequencies were lower than the boundary and licking rightward when the tone frequencies were higher than the boundary. Imaging experiments were started after 2-5 days of training in the second training stage.

For the rule-switching flexible categorization (RSFC) task, the training procedure was described previously^36^. In brief, the behavior training consisted of four stages. The first stage is the two tones discrimination task (7 kHz and 28 kHz or 4 kHz and 16 kHz), reaching a criterion of > 85% correct rate. The second training stage is the reversal shaping, in which there are two different types of trial blocks alternating within a session. In one type of block, each trial presents either 7 kHz (choosing left) or 14 kHz (choosing right). In the other type of block, each trial presents either 14 kHz (choosing left) or 28 kHz (choosing right). Thus, the choice for 14 kHz tone is reversed in alternating blocks of trials, while the choice for 7 kHz or 28 kHz remains unchanged across blocks. The block switch occurs without any explicit cues once the correct rate of 14 kHz trials reached 75%. Once the performance in the 14 kHz trials reaches 80% correct for 3 consecutive days, the third training stage starts. The third stage is the one-tone reversing stage, with each block of trials containing 3 trial types (7 kHz, 14 kHz, or 28 kHz). The choice for 14 kHz needs to be reversed in alternating blocks, while the choice for 7 kHz or 28 kHz remains unchanged. The final stage is the full version of the flexible categorization task, which includes 7 tone stimuli (7 kHz, 9 kHz, 11 kHz, 14 kHz, 18 kHz, 22 kHz, 28 kHz). In alternating blocks of trials, the choice contingencies for the 3 intermediate frequencies (11 kHz, 14 kHz and 18 kHz, reversing stimuli) are reversed across blocks, but remain unchanged for the two lower (7 kHz and 9 kHz) and higher (22 kHz and 28 kHz) tones (non-reversing stimuli). The task design effectively sets two alternating categorization rules (rule 1, with category boundary at 19 kHz; rule 2, with category boundary at 10 kHz) in different blocks of trials.

For all the blocks, the trial number and reward size are balanced for the left and right choice sides. To minimize the difference in distribution of tone frequencies in different block types, while maintaining balanced choice values, the occurrence of different sound stimuli is similar in left or right trials, i.e., ∼25% respectively for 7 kHz or 9 kHz (left trials), ∼10% respectively for 11 kHz, 14 kHz, 17 kHz, 21 kHz or 28 kHz (right trials), and in high boundary blocks, ∼25% for 21 kHz or 28 kHz (right trials), ∼10% for 7 kHz, 9 kHz, 11 kHz, 14 kHz or 17 kHz (left trials). Normally mice performed 450-650 trials per session/day, providing sufficient statistic power.

To quantify the categorization behavior, the psychometric function was obtained by plotting the proportion of choices against the tone frequency, fitted with a 4-parameter sigmoidal (logistic) function:

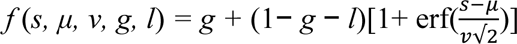

where f is the probability that the animal would make a right choice, *s* is the tone frequency in the octave, *μ* is the mean value of the distribution representing the subjective category boundary (point of subjective equality, PSE), *v* is the variation in the distribution representing the subject’s discrimination sensitivity, and *g* and *l* are the guess rate and lapse rate, respectively. And erf is the error function.

The BSRL model capturing the inference-based flexible categorization behavior has been described in our previous study^36^. Briefly, in this model, the perceived stimulus, ŝ_t_, is normally distributed with constant variance σ^2^ around the true stimulus frequency:

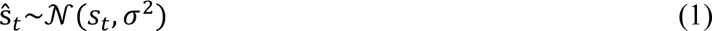

Given ŝ_t_, the agent has the belief that the stimulus belongs to the high- or low-frequency category according to two possible boundaries:

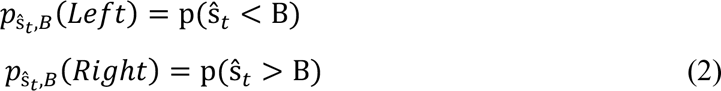

The expected values of left and right choices (A_t_(*i*)) are computed as following where ω_B_ is the subjective estimate of the category boundary:

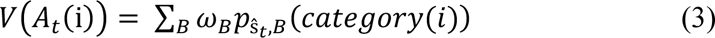

The values drive choices via a softmax function:

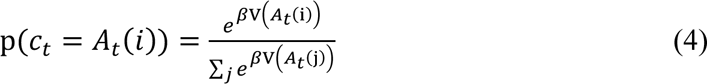

Where β is an inverse temperature parameter which will be fit to data.

The ω_B_ is updated by incorporating the value prediction error δ_t_ as:

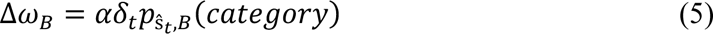

Where the value prediction error is:

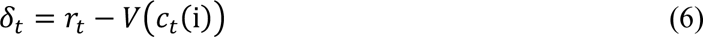

This model contained 4 parameters including *σ*, β, α, and initial value of ω_B_. We used the negative log likelihood (NLL) as cost function, and *fmincon* function from Matlab to minimize the NLL. The model was fit separately for individual mouse, using threefold cross-validation. Additional model testing and the comparison of the performance of BSRL model with a standard RL model in capturing the behavioral data were carried out in our previous study^36^.

### Two-photon calcium imaging

Calcium imaging was performed using a custom-built two-photon microscope (MIMMS 1.0, Janelia Research Campus) with minimized scanning noise in a sound proofing system reducing ambient noise, as described previously^36^. GCaMP6s was excited using a Ti-Sapphire laser (Chameleon Ultra II, Coherent) tuned to 920 nm, with average power at ∼70 mW measured at the entrance pupil of the objective. Images were acquired using a 16x 0.8 NA objective (Nikon), and the GCaMP6s fluorescence was isolated using a bandpass filter (525/50, Semrock) and detected using GaAsP photomultiplier tubes (PMT, 11706-40, Hamamatsu). Horizontal scanning was through a resonant galvanometer (Thorlabs; 8 kHz line rate, bidirectional). The imaging frame rate is ∼30 Hz with 512 × 512 pixels. Soma and dendrites from the same region were imaged with different focal depth in different behavioral sessions with randomized order. To avoid potential repeated sampling, different imaging fields were used in different sessions based on the localization using blood vessels and images from somatic imaging fields. For each mouse, 2-3 locations were imaged at different focal depths. The field of view was 300 × 300 µm for soma imaging, ∼ 250 × 250 µm for trunk imaging, and ∼100 × 100 µm for tuft imaging, with respective focal depths as shown in Extended Data Fig. 8. The acquisition system was controlled using ScanImage. For each animal, the optical axis was adjusted (45-50 degrees from vertical) to be perpendicular to the imaging window in the auditory cortex. Different fields of view in the same mouse were imaged on different sessions. At the initiation of each trial, ScanImage acquisition was triggered by a starting signal from the PX-Behavior System, continuous scanning covering the entire behavioral trial were recorded.

For 2-plane calcium imaging from the dendrites and somas, a piezo objective positioner kit (P-725K129, Physik Instruments, Germany) was used to drive the objective to acquire images with at two focal planes separated by a distance of 40 to 380 μm with alternating frames, at 6.9∼8.7 Hz per plane. The field of view was 300 × 300 µm, with 512 ×512 pixels. The signal correlation for z-scored activity was calculated between images from the two planes for each ROI. The dendritic and somatic ROIs from the same neurons were first identified based on the signal correlation between images from the two focal planes and confirmed visually based on z-stacks acquired with a 1-μm z step between the two imaging planes^62^.

### Optogenetic inhibition of dendritic activity

To optically activate the SST neuron while simultaneously imaging the L5 ET neurons dendrites, red light (635 nm) was delivered using a diode laser (Shanghai Laser & Optics Century Co., Ltd.) to stimulate ChrimsonR in the same region of two-photon imaging in ACx. The 635 nm laser was delivered through the objective via a custom-designed light path with an extra dichroic mirror (FF705-Di01, Semrock) and a modified primary dichroic mirror (FF594-Di04, Semrock) before the entrance pupil of the objective. The modified primary dichroic mirror passes the 635-nm laser into the objective and the reflected GCaMP6s emission light to the detection arm. The 635-nm light coming out of the objective was ∼1 mm in diameter at the focal plane, with a total power of 1.2-2.4 mW. To prevent optical artifacts from optical stimulation during imaging, we used a custom-designed Optogenetic Controller for 2P to control the PMT shutter time and the red-light delivery time relative to the 2P scanning phase. The red-light was delivered only in the fly-back period of the resonant scanner during each line scan, during which the PMT was turned off by the shutter circuit to protect the photo detector. The red light was illuminated with a duration of ∼31 µs in each line period, and with a repetition rate of ∼8 kHz (determined by the resonant scanner of the 2P microscope). In each photostimulation trial, the onset of red-light stimulation was 500 ms before sound onset and lasted for 4 s. In each session, the proportions of photo stimulation trials were ∼50%, which were randomly interleaved with control trials.

### Imaging data processing

Custom-written software based on MATLAB^20^ or Suite2p^76^ was used for imaging data processing. We performed frame-by-frame registration to correct brain motion in the x-y direction. Z-axis movement was visually inspected and imaging sessions with significant z-axis movement were discarded. For the motion correction registration, the imaging frames from each session were aligned to a target image frame using a cross-correlation-based registration algorithm (discrete Fourier transformation, DFT). The target image was obtained by taking the mean projection of visually identified frames with few motion artifacts. For relatively sparse trunk fields, the target image was averaged from several frames from different trials to reduce misalignment for “star-like” targets. To extract fluorescence signals from individual neurons, regions of interest (ROIs) were manually drawn based on the neuronal shape using a custom GUI software in MATLAB^20^, or using the Suite2p^76^. Mean, maximum intensity and standard deviation values of all frames of a session were used to determine the boundaries of the ROIs.

The pixel intensity values within each ROI in a frame were averaged as the fluorescence intensity of a ROI at that frame time. The fluorescence intensity of each ROI was then extracted over all the frames as the fluorescence time series (*F*). Before calculating the ΔF/F0, slow calcium fluorescence changes were removed by determining the distribution of fluorescence values in a 20-second interval around each sample time point and subtracting the 8th percentile value. For each ROI, *ΔF/F0* (%) was calculated as *(ΔF/F0*) × 100, where *F*0 is the index of the peak of the histogram of *F*. The *ΔF/F0* traces were used for all subsequent analyses. Missed trials were excluded from all analyses.

### Quantification and statistical analysis of neuronal activity

In part of our analysis, we used n-way ANOVA to determine whether an ROI shows significant coding (*p* < 0.05) for a given behavioral variable (Extended Data Fig. 2; Extended Data Fig. 9). For comparing the contributions to an ROI activity across multiple variables, we used ANOVA with multiple comparisons (Extended Data Fig. 6f-i).

We used a receiver-operating characteristic (ROC) analysis to measure discrimination ability for the choice or rule of soma, trunk, and tuft activity. The area under the ROC curve (auROC) was used as the discriminability measure for a single ROI. To construct the ROC curve, we used the average calcium signals from a 1 s window around the peak response as the activity for the ROI in each trial. To determine the statistical significance of single ROI discrimination, a null ROC value distribution was computed by randomly assigning the neuronal responses to the corresponding task variables. This process was repeated 1000 times to obtain a null distribution of auROC values. The threshold for significance was defined as the 95th percentile of the shuffled auROC values.

The selectivity index (SI) for rules or choices was calculated based on the auROC value as:

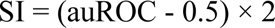

This index directly reflects the auROC value with the sign corresponding to the direction of selectivity. For rule selectivity, an index of 1 indicates perfect selectivity for rule 1, −1 for rule 2, and 0 for no selectivity. The SI for rule was calculated using non-reversing correct trials in the preferred side. To calculate the auROC value for rule discrimination by pre-stimulus activity, we use 1 s period before stimulus onset.

To examine the simultaneous contribution of different task variables to the neuronal activity, we modeled the dendritic and somatic activity using a generalized linear model (GLM) approach^24^.

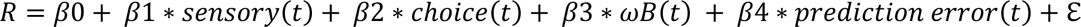

In each trial, *t*, various variables such as sensory, choice, ω_B_ (rule estimate), and prediction error were used as predictors to predict the neuronal activity of an ROI. For sensory, the variable is a categorical with different frequencies labeled from 1 to 7. Choice is a binary variable labeled as 1 for left choice and 2 for right choice. Rule estimate was also labeled as 1 for the low boundary block and 2 for the high boundary block or using boundary estimation value from BSRL-model. Prediction error value was calculated from BSRL-model. To model the contribution of different task variables at different time epochs, the calcium signals from three different time epochs were used: epoch 1, calcium signals averaged from a 1 s period after stimulus onset; epoch 2, averaged calcium signals from the 2 s period after the first choice lick following stimuli onset; epoch 3, averaged calcium signals from the 2 s period of answer time.

A full model with all predictors provided the explained variance *R_full_*^2^ for each ROI. ROIs with *R_full_*^2^ > 0.1 were included for the further GLM analysis, unless stated otherwise. The relative contribution of each variable to the explained variance F was evaluated as:

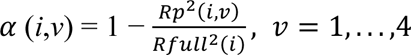

where *Rp*^2^(*i*,*v*) is the explained variance by the partial model without the variable *v*.

To quantify the changes in the encoding for different variables following dendritic inhibition (Fig. 5), neurons with > 5% variance explained by the full regression model were included, and the effects were separately assessed for choice, sensory and rule selective neurons with >10% relative contribution to the explained variance from each variable.

For population decoding (Extended Data Fig. 2p, q), we trained a Support Vector Machine (SVM) classifier with a linear kernel to predict choices using simultaneously recorded neurons. In each session, the data were arranged into an M-by-N matrix, where M represented the number of trials and N represented the number of ROIs. Each element in the matrix represents the responses of an ROI in each trial. For decoding with activity from different trial time epoch, the responses were averaged from different time windows (0.8 s): pre-sound, from stimulus onset, and after answer choice, respectively. Leave-one-out cross-validation was used to calculate the decoding accuracy, and this process was repeated 100 times to obtain the average decoding accuracy. The neuronal activity in trials from the L5 IT and ET neurons was normalized separately, and the number of ROIs from different cell types was balanced. Only correct trials were used, and the number of trials for training was the same in both cell types.

We also used mutual information (MI)^52,77^ to quantify the information conveyed by neuron activity in relation to behavioral variables: sensory stimulus, choice, and rule. We use the k-nearest neighbor (KNN) method^78^ to calculate MI, which is based on entropy estimates from KNN distances (k = 5). To assess significance, the MI value for each ROI was compared with the mean MI value from a shuffled dataset (comprising 500 shuffles); MI values exceeding 2 standard deviations above the average of shuffled MI were considered statistically significant.

The statistical comparisons were conducted using MATLAB (MathWorks). The student’s t-test, two-sample t-test, and paired-sample t-test were used for normally distributed data, while the Wilcoxon signed-rank test and Wilcoxon rank-sum test were used for non-parametric data. n.s. indicates that the results were not significant (p > 0.05), while *, **, and *** denote p values of < 0.05, < 0.01, and < 0.001, respectively. The standard error of the mean (SEM) is represented by error bars or shaded regions surrounding line plots, unless otherwise stated. Sample sizes were not predetermined using statistical methods, but they are comparable to those reported in previous publications^36,38,48^. The data collection and analysis were not blinded except for the tracing performed in single-neuron reconstruction, as all behavioral variables were objectively measured by an automated hardware and software system that did not require human intervention.

### Chemogenetic silencing of different cell types

We used a designer receptor exclusively activated by designer drugs (DREADD)-based method to silence the L5 IT and ET neurons in the auditory cortex, respectively (Extended Data Fig. 3). The expression of hM4D(Gi) in different cell types is as described in the previous sections. The Clozapine-N-Oxide (CNO, Sigma) was dissolved in a saline solution (0.9% NaCl) to make a stock solution of 20 mg/ml, which was stored at −20°C. On each day of the experiment, the stock solution was diluted to a working concentration of 0.2 mg/ml. CNO was administered at a dose of 2 mg/kg of mice body weight (0.2 mg/ml) through intraperitoneal (i.p.) injection 30-40 minutes before behavioral testing. Saline with same dose was used as control.

### Histology

After performing imaging and manipulation, the mice were given deep anesthesia and underwent transcranial perfusion with a solution of 0.9% NaCl, followed by a 4% paraformaldehyde (PFA) in PBS solution. The brains were then post-fixed in the same PFA solution overnight and dehydrated using 30% sucrose. Coronal brain sections of 50-80 µm were cut using a cryostat (Leica CM1950) and imaged on a virtual slide microscope (Olympus VS120).

### Single-neuron reconstruction

The axon and dendrite of single neuron were labeled, imaged and reconstructed as described previously^43^. Neurons in the auditory cortex were sparsely labeled using a mixture of a diluted virus expressing Cre-recombinase and a Cre-recombinase dependent reporter virus. The full volume of each brain was imaged using fluorescence micro-optical sectioning tomography (fMOST)^79^. Individual neurons were reconstructed manually using FNT. To ensure the validity of tracing result, every neuron was traced by two independent human tracers. The results were then merged by a third human tracer. Each brain sample was registered to the Allen Common Coordinate Framework (CCFv3) using rigid and non-rigid registration, which was accomplished by using Computational Morphometry Toolkit software. At last, reconstructed neurons were mapped onto CCFv3 to determine the brain area associated with each axonal node.

## Extended Data

**Extended Data Fig. 1.**
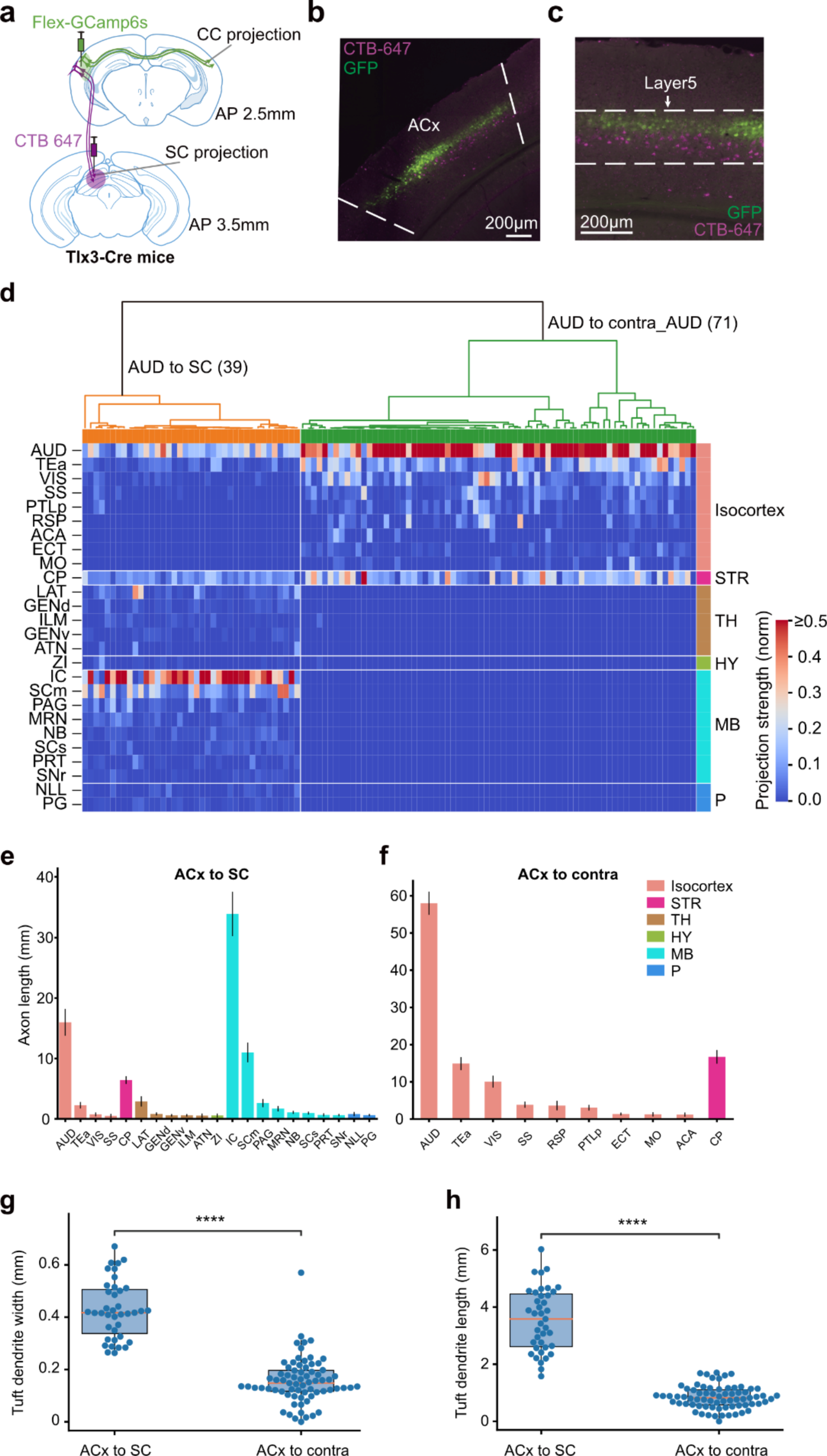
Projection-defined ACx L5 neuron types. **a**, Schematic showing concurrent labeling of SC-projecting and contra-projecting ACx L5 neurons. SC-projecting neurons were retrogradely labeled by injecting cholera toxin subunit B (CTB)-647 into the ipsilateral SC. Contra-projecting ACx L5 neurons were labeled by injecting AAV-Syn-FLEX-EGFP in the contralateral auditory cortex of the Tlx3-cre mouse. **b** and **c**, Soma of the two projection cell types were largely non-overlapping. **d**, Clustering analysis of the two projection cell types based on their whole-brain projection targets. Data are from whole brain reconstruction of single ACx neurons. **e**, SC-projecting neurons show stronger projections to TH, HY, MB, and P. TH: thalamus, HY: hypothalamus, MB: midbrain, P: pons. Error bar, mean ± SEM. **f**, The contra-projecting ACx L5 neurons mainly project to the Isocortex and striatum. **g** and **h**, Box plots showing the width and total length of the distal dendritic tuft of SC-projecting and contra-projecting ACx L5 neurons. Two-sample *t*-test, ****, p < 0.0001.

**Extended Data Fig. 2.**
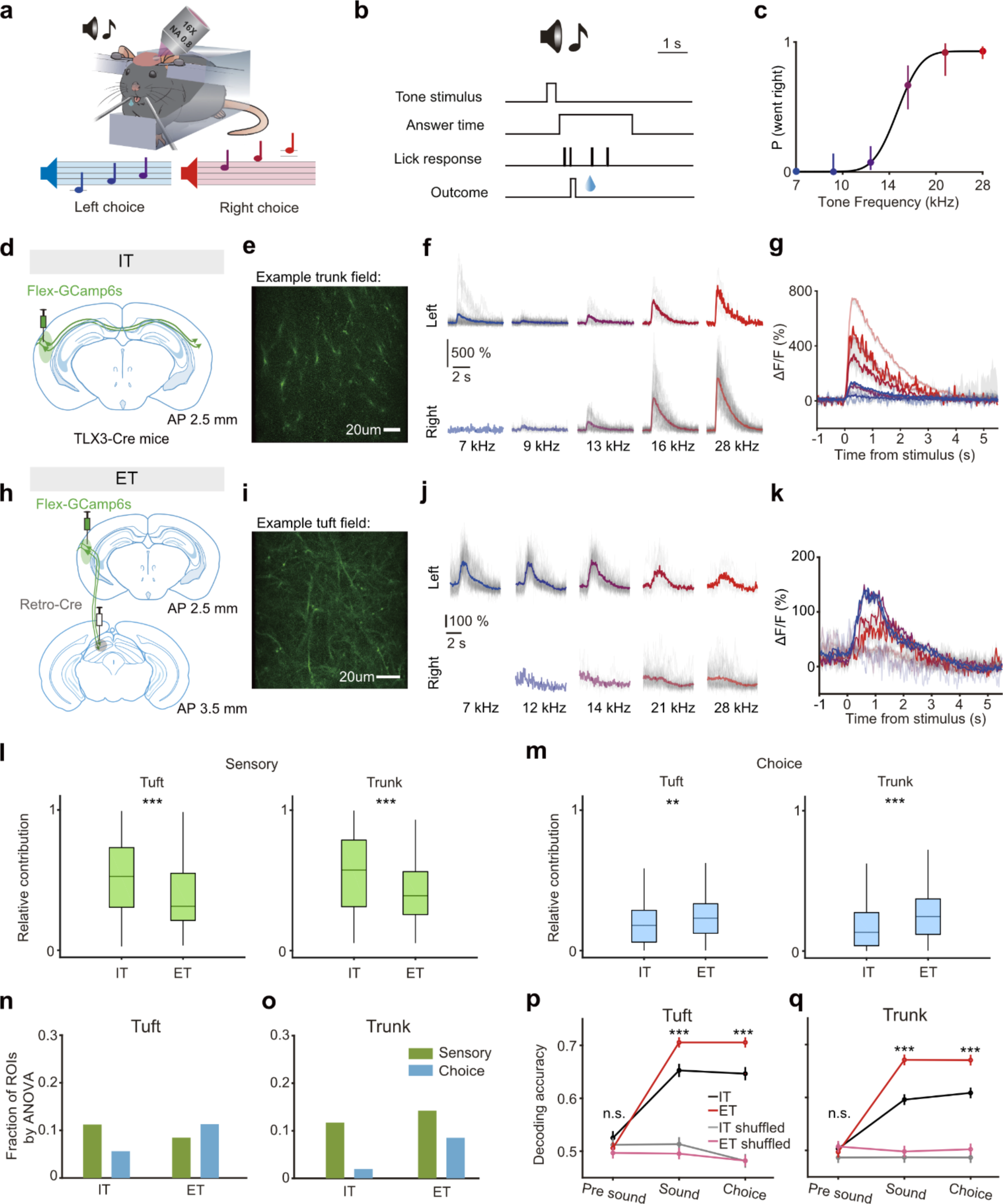
Different coding properties of ET and IT L5 neurons. **a**, Schematic showing *in vivo* two-photon imaging of auditory cortex during auditory guided 2AFC task. **b**, Time structure for each trial. **c**, Example psychometric function of one behavior session. Error bars represent 95% confidence intervals. **d**, Schematic showing labeling of IT neurons in the ACx with GCamp6s via virus injection in Tlx3-Cre mice. **e**, Example imaging field of the Ca^2+^ indicator GCaMP6s in layer 5 IT dendrites. **f**, Example traces of Ca^2+^ signals from an example sensory selective dendritic ROI of IT neurons, with separate groups of trials presenting different tone frequencies (colors) with different choices (upper and lower), gray lines showing individual trial activity. **g**, Mean traces from **f**, shadow indicate the SEM. **h** to **k**, labeling and Ca^2+^ imaging of SC-projecting ET neurons, similar as in **d-g**. **l** and **m**, Relative contribution of sensory (**l**) and choice (**m**) of tuft and trunk ROIs in IT and ET neurons, based on GLM modeling. Sensory, green (**l**); choice, blue (**m**). Wilcoxon rank-sum test, ***, p < 0.001, **, p < 0.01. **n**, Fraction of sensory and choice selective *tuft* ROIs in IT and ET neurons, calculated by ANOVA. IT neurons, sensory selective 11%, choice selective 5%, n = 1050 ROIs; ET neurons, sensory selective 8%, choice selective 11%, n = 1026 ROIs. **o**, Fraction of sensory and choice selective *trunk* ROIs in IT and ET neurons, calculated by ANOVA. IT neurons, sensory selective 11%, choice selective 1%, n = 913 ROIs; ET neurons, sensory selective 15%, choice selective 9%, n = 492 ROIs. **p**, Population decoding accuracy at different trial epochs (pre-sound period 800 ms, sound period 800 ms, and choice period 800 ms) for *tuft* ROIs in IT and ET neurons, summarized for all sessions. Red lines, ET neurons, 11 sessions. Black lines, IT neurons, 8 sessions. Decoding accuracy for ET neurons, sound period, 0.71 ± 0.01, choice period, 0.71 ± 0.01; IT neurons, sound period, 0.65± 0.01, choice period, 0.65 ± 0.01 (two-sample *t* test, ***, p < 0.001; n.s., not significant). Error bars indicate SEM. **q**, Decoding accuracy as in **p** for trunk ROIs of IT and ET neurons. IT neurons, sound period, 0.60 ± 0.01; choice period, 0.6 ± 0.01; n = 13 sessions. ET neurons, sound period, 0.67 ± 0.01; choice period, 0.67 ± 0.01(two-sample *t* test, ***, p < 0.001; n.s., not significant). Error bars indicate SEM.

**Extended Data Fig. 3.**
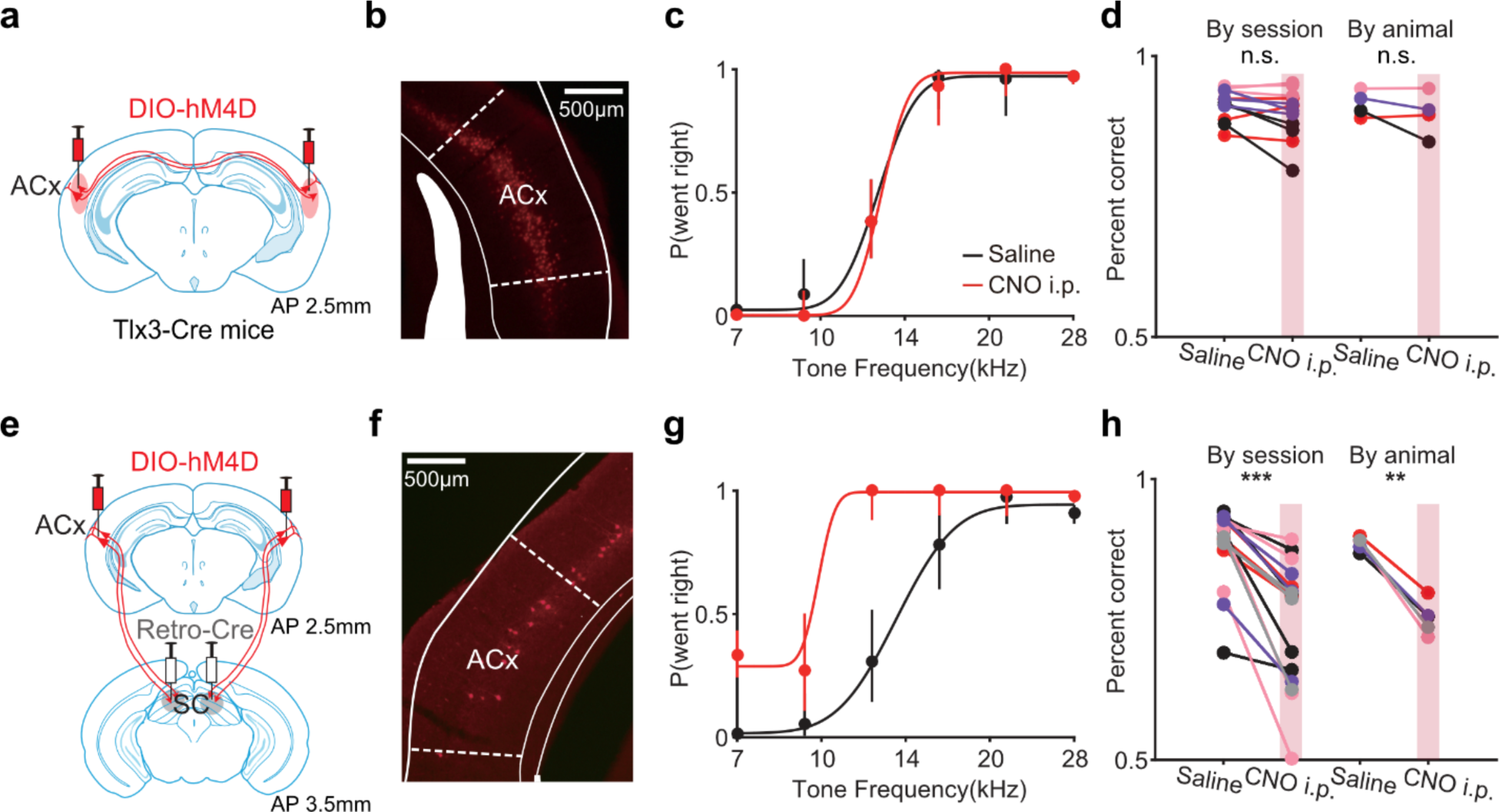
Causal requirements of the ET and IT neurons in auditory for perceptual decision behavior. **a**, Schematic showing virus injection of AAV-DIO-hM4D in IT neurons in bilateral ACx using Tlx3-Cre mice. **b**, Example histological image showing expression of hM4D in ACx IT neurons. **c**, Psychometric functions showing example behavioral performance with saline (black) or CNO injection (red) in two consecutive sessions. Error bars represent 95% confidence intervals. **d**, Summary of behavioral performance from all sessions and animals, compared between with and without IT neuron silencing. By session (left), p = 0.09, n = 12 session pairs, Wilcoxon sign-rank test, n.s., not significant. By animal (right), p = 0.89, n = 4 mice, Wilcoxon rank-sum test, n.s., not significant. **e**, Schematic showing virus injection of AAV-DIO-hM4D in ET neurons in bilateral ACx with Retro-Cre injected in bilateral SC. **f**, Example histological image showing expression of hM4D in ET neurons in ACx. **g**, Example psychometric functions as in **c** for ET neurons in ACx. **h**, Summary of behavioral performance from all sessions and animals, compared between with and without ET neuron silencing. By session (left), p = 2.93 × 10^-4^, n = 17 session pairs, Wilcoxon sign-rank test, ***, p < 0.001. By animal (right), p = 0.79 × 10^-2^, n = 5 mice, Wilcoxon rank-sum test, **, p < 0.01.

**Extended Data Fig. 4.**
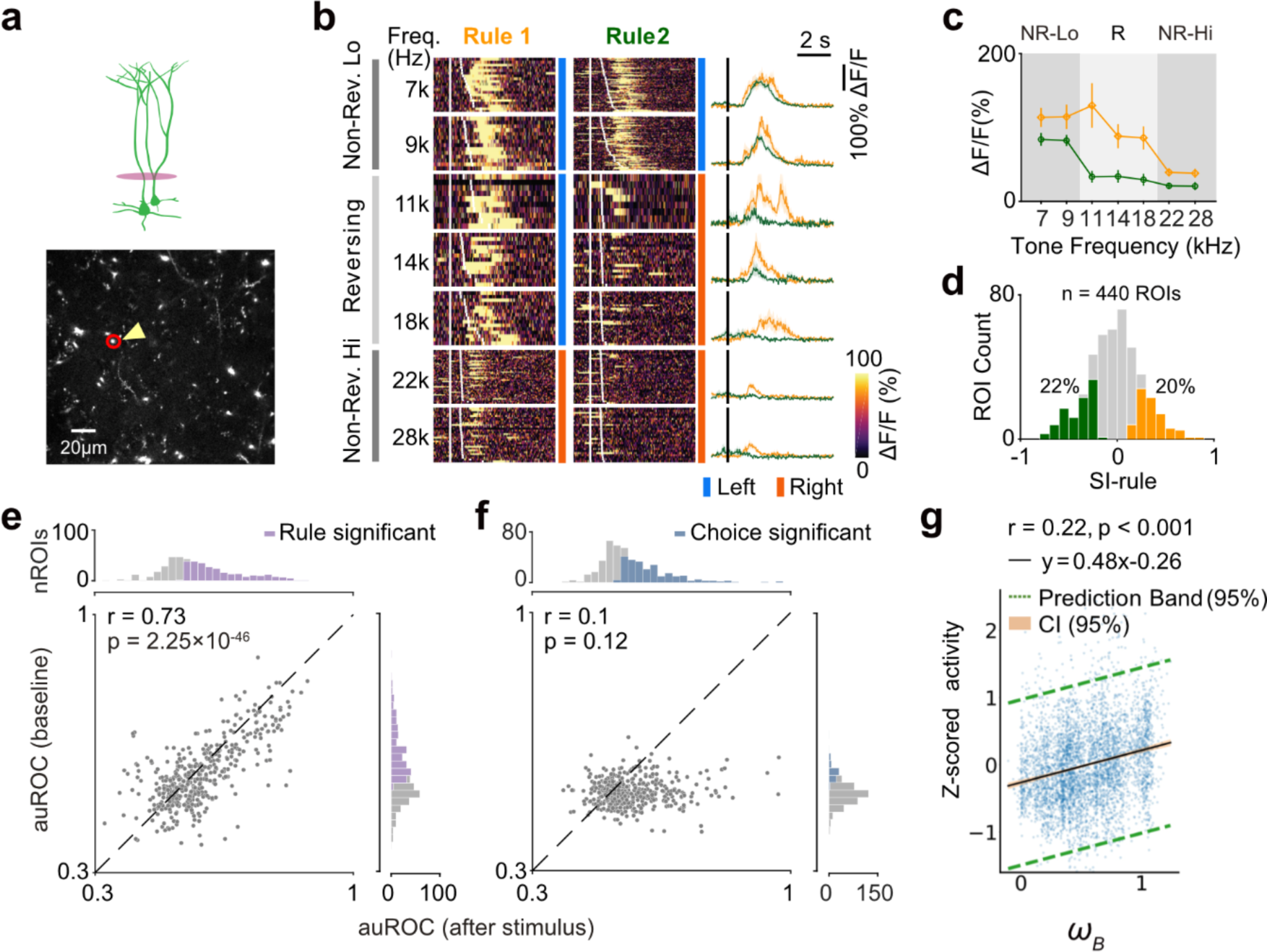
Rule information coding in dendritic trunks during rule-switching flexible categorization. **a**, Example two-photon image of dendritic trunks of ET neurons in ACx. **b**, Color plot of trial-by-trial Ca^2+^ signals from the trunk ROI in **a** with red circle, showing rule selectivity in an example behavioral session. As in Fig. 2b. **c**, Summary of the Ca^2+^ signals in **b**. **d**, Distribution of rule selectivity index for all trunk ROIs (n = 440 ROIs from 5 mice). **e**, Correlation of auROC values for rule discrimination before and after stimulus onset in trunk ROIs. Each dot represents auROC values from one ROI. X-axis, auROC value computed from peak 1 s activity after stimulus onset; y-axis, auROC value computed from 1 s activity before stimulus onset. Histograms on the top and right represent the distribution of auROC values with color indicating ROIs with significant rule selectivity (p < 0.05, permutation test, n = 1000 random permutations). **f**, Correlation of auROC values for choice discrimination before and after stimulus similar as in **e**. **g**, Trial-by-trial activity from rule selective trunk ROIs as a function of the rule estimate values calculated from the BSRL model (4713 trials from 183 rule selective ROIs).

**Extended Data Fig. 5.**
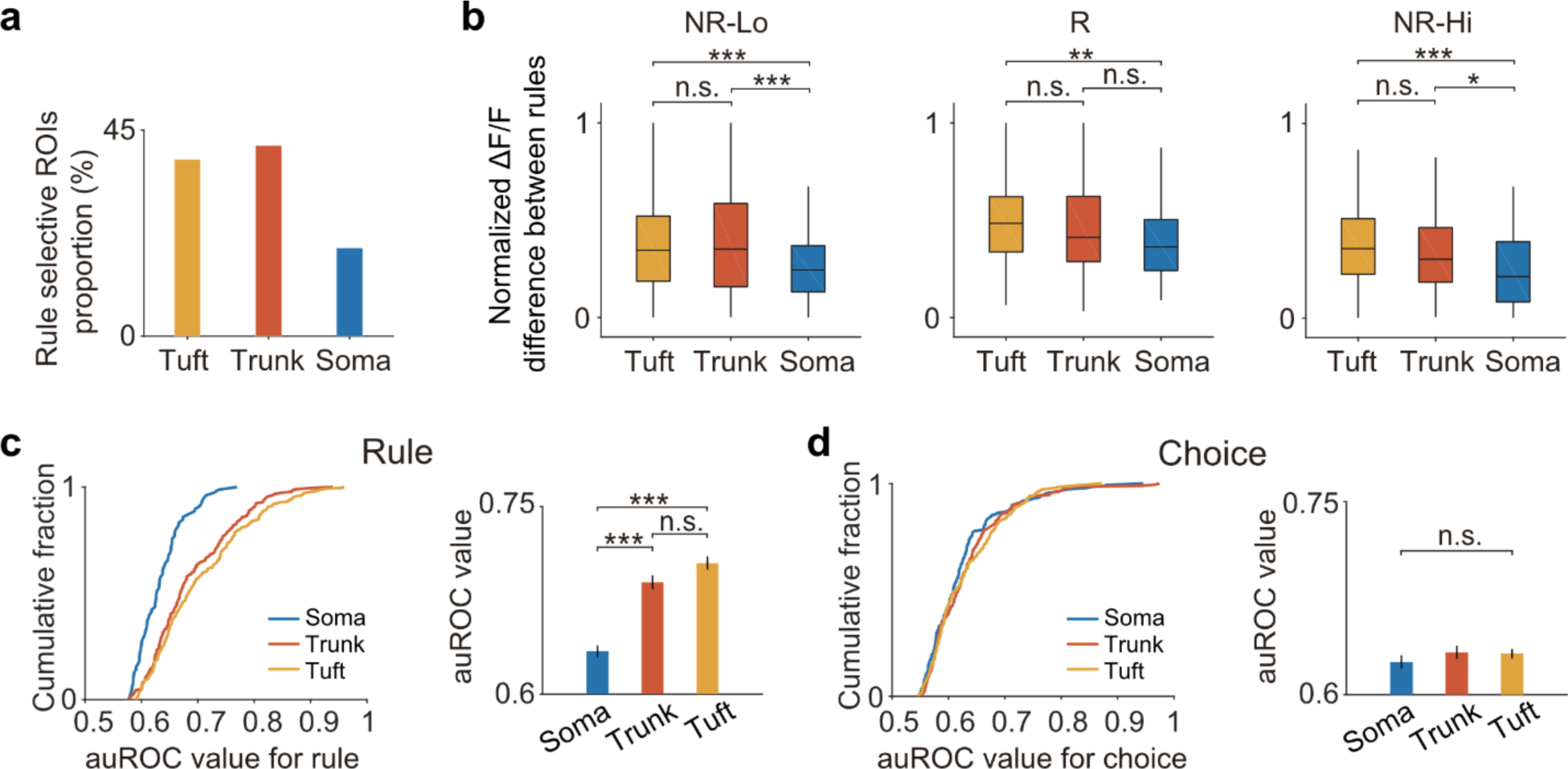
Rule and choice discrimination by different subcellular compartments. **a**, Proportion of rule selective ROIs revealed by auROC analysis. Soma (blue, n = 78/405), trunk (orange, n = 183/440), and tuft (yellow, n = 249/645). **b**, Comparison of normalized Ca^2+^ activity (each ROI was normalized by its max activity) difference between rules for 3 compartments under non-reversing low-frequency trials, reversing trials, and non-reversing high-frequency trials respectively. NR-Lo, Tuft-trunk, p = 0.91, Soma-trunk, p = 2.36 × 10^-4^, Soma-tuft, p = 4.21 × 10^-4^; R, Tuft-trunk, p = 0.25, Soma-trunk, p = 0.17, Soma-tuft, p = 6.26 × 10^-3^; NR-Hi, Tuft-trunk, p = 0.27, Soma-trunk, p = 1.75 × 10^-2^, Soma-tuft, p = 1.81× 10^-4^, ANOVA with multiple comparisons, ***, p < 0.001; **, p < 0.01; *, p < 0.05; n.s., not significant. Soma (blue, n = 78), trunk (orange, n = 183), and tuft (yellow, n = 249). **c**, Comparison of auROC values for rule discrimination in all rule selective ROIs between soma (blue, n = 78), trunk (orange, n = 183), and tuft (yellow, n = 249) compartments. Right, averaged auROC values. Error bars indicate SEM. Tuft-trunk, p = 0.09, Soma-trunk, p = 2.15 × 10^-7^, Soma-tuft, p = 9.58 × 10^-10^, ANOVA with multiple comparisons, ***, p < 0.001; n.s., not significant. **d**, Comparison of auROC values for choice discrimination in all choice selective ROIs between soma (blue, n = 188), trunk (orange, n = 205), and tuft (yellow, n = 313). Right, averaged auROC values. Error bars indicate SEM. ANOVA, F = 0.71, p = 0.49, n.s., not significant.

**Extended Data Fig. 6.**
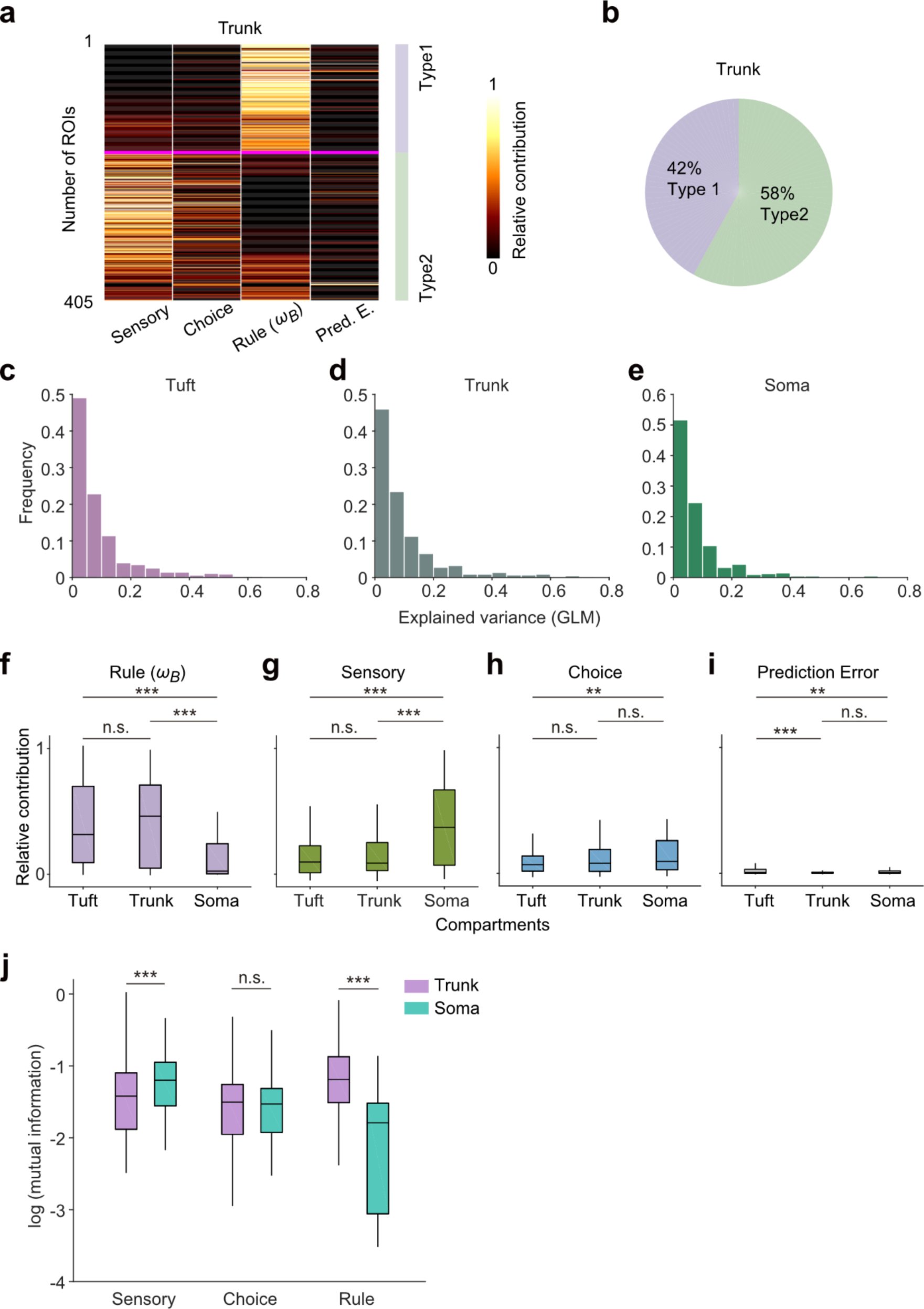
Contribution of task variables to Ca^2+^ signals in subcellular compartments. **a**, Clustering of dendritic trunk ROIs based on relative contributions by different task variables to the explained variance of trunk activity. Type 1 and type 2 ROIs are the primary clusters. Clusters were sorted by distance calculated using the hierarchical-linkage-cosine method. **b**, Proportion of each cluster type in trunk. **c** to **e**, Distribution of the explained variance by the full model of GLM for the activity in ROIs from different subcellular compartments. Tuft (**c**), 645 ROIs; trunk (**d**), 440 ROIs; soma (**e**), 405 ROIs. **f**, Summary of relative contributions of rule estimate to activity in tuft, trunk, and soma ROIs with explained variance > 0.1, ANOVA with multiple comparisons, ***, p < 0.001; n.s., not significant. **g**, Similar as in **f**, for the contribution of the sensory variable, ANOVA with multiple comparisons, ***, p < 0.001; n.s., not significant. **h**, Similar as in **f**, for the contribution of choice, ANOVA with multiple comparisons, **, p < 0.01; n.s., not significant. **i**, Similar as in **f**, for the contribution of the prediction error. ANOVA with multiple comparisons, **, p < 0.01; n.s., not significant. **j**, Summary of mutual information (MI) in trunk (n = 130) and soma (n = 93) for ROIs with explained variance > 0.1. MIs are compared between trunk and soma for sensory (p = 0.0027), choice (p = 0.81), and rule (p = 1.20 × 10^-14^). Wilcoxon rank-sum test, ***, p < 0.001; n.s., not significant.

**Extended Data Fig. 7.**
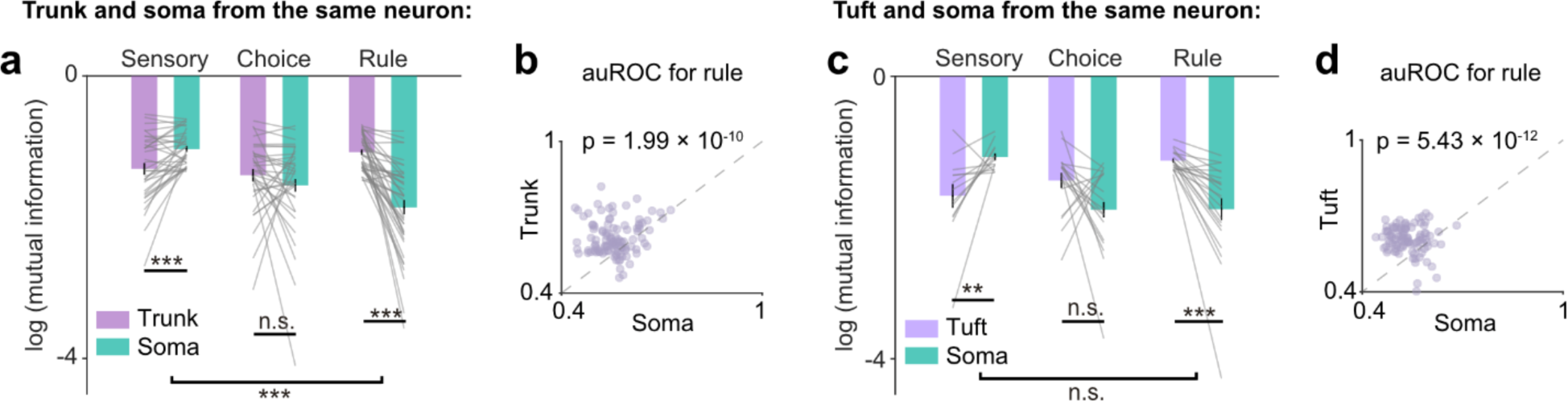
Task variable coding by sub-neuronal compartments within single neurons. **a**, Mutual information (MI > 0) compared between soma and trunk within the same individual neurons for sensory stimuli (MI-sensory) (p = 4.72 × 10^-4^, Wilcoxon sign-rank test, n = 34 neurons with significant somatic MI-sensory); for choices (MI-choice) (p = 0.23, Wilcoxon sign-rank test, n = 34 neurons with significant MI-choice); for rules (MI-rule) (p = 3.78 × 10^-8^, Wilcoxon sign-rank test, n = 41 neurons with significant trunk MI-rule). Error bars represent SEM. Difference between sensory change and rule change, Wilcoxon rank-sum test, p = 6.47 × 10^-4^. ***, p < 0.001; n.s., not significant. **b**, auROC value for significant rule coding ROIs from same neuron (soma and trunk), p = 1.99 × 10^-10^, Wilcoxon sign-rank test, n = 109. **c**, Mutual information (MI > 0) compared between soma and tuft within the same individual neurons for sensory stimuli (MI-sensory) (p = 4.63 × 10^-3^, Wilcoxon sign-rank test, n = 13 neurons with significant and positive somatic MI-sensory); for choices (MI-choice) (p = 0.1, Wilcoxon sign-rank test, n = 18 neurons with significant MI-choice); for rules (MI-rule) (p = 7.98 × 10^-5^, Wilcoxon sign-rank test, n = 21 neurons with significant tuft MI-rule). Error bars represent SEM. Difference between sensory change and rule change, Wilcoxon rank-sum test, p = 0.67. ***, p < 0.001; **, p < 0.01; n.s., not significant. **d**, auROC value for significant rule coding ROIs from same neuron (soma and tuft), p = 5.43 × 10^-12^, Wilcoxon sign-rank test, n = 100.

**Extended Data Fig. 8.**
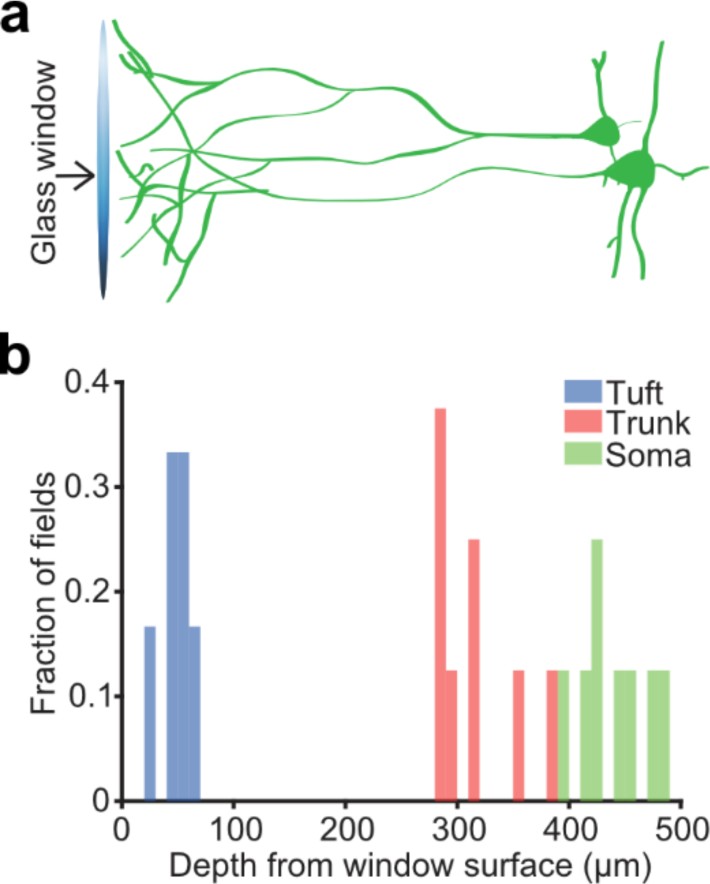
Imaging field depth information. **a**, Schematic illustrating the range of ET neurons dendritic and somatic positions relative to the scale shown in **b**. **b**, Distribution of imaging fields over the focal depth under imaging window corresponding to the subcellular locations in **a**.

**Extended Data Fig. 9.**
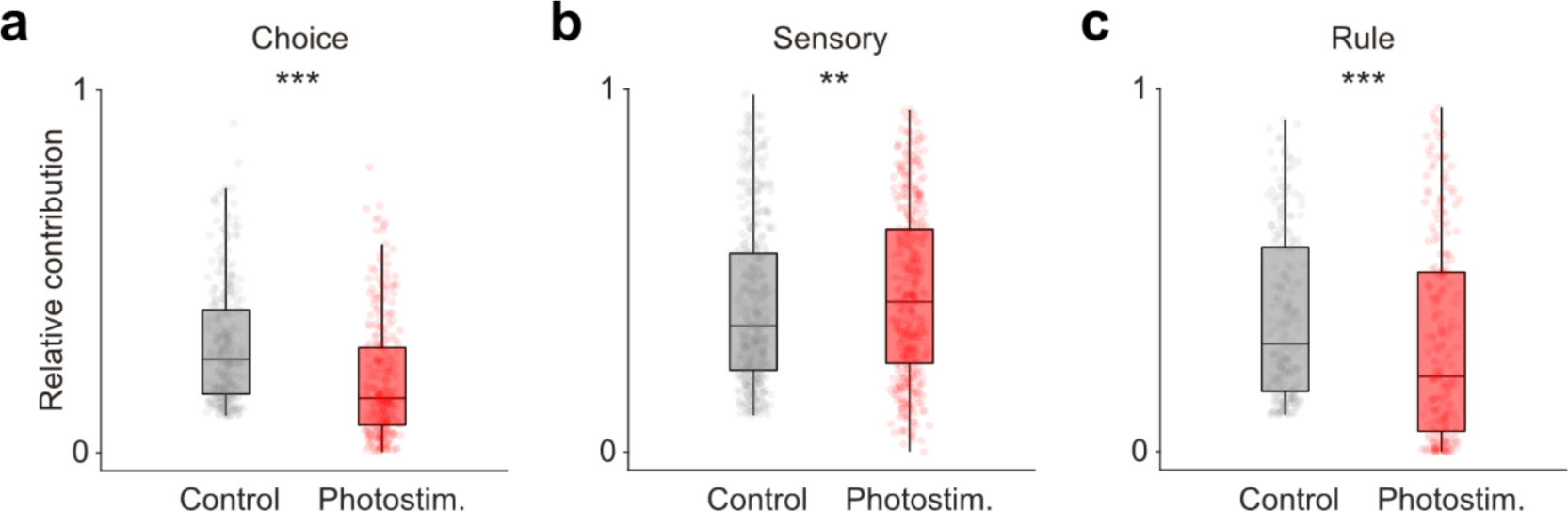
Relative contribution of task variables to dendrites Ca^2+^ activity with and without photostimulation. Relative contribution to the Ca^2+^ signals in dendrites compared between control and photostimulation trials, as in Fig. 5. For choice selective ROIs, p = 2.99 × 10^-20^, n = 295, (**a**); sensory selective ROIs, p = 1.8 × 10^-3^, n = 419, (**b**); rule selective ROIs, p = 1.29 × 10^-6^, n = 227, (**c**). Wilcoxon signed-rank test, **, p < 0.01; ***, p < 0.001.

**Extended Data Table 1.**
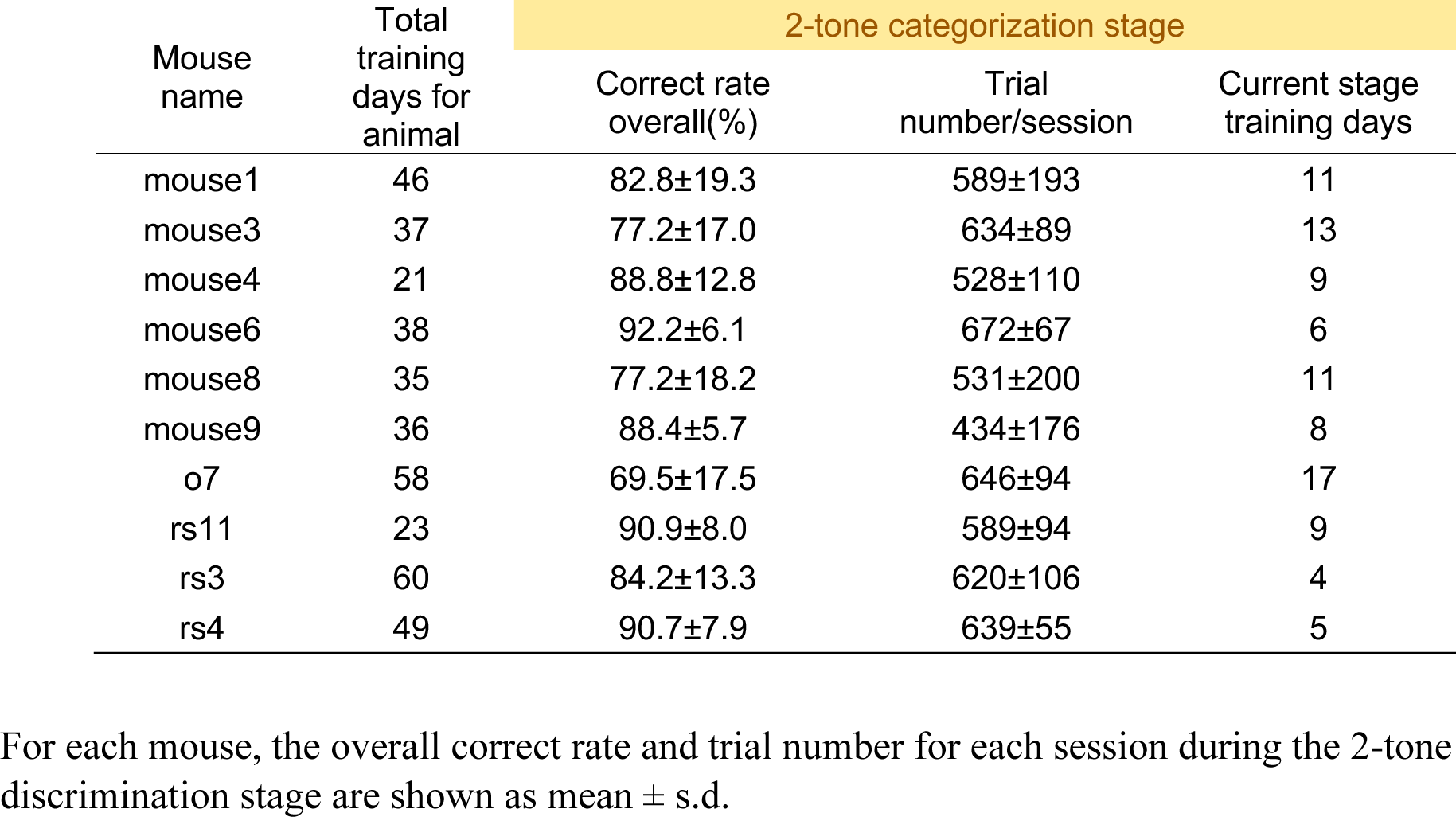
Behavior training data for the 2-tone categorization stage.

| Mouse name | Total training days for animal | 2-tone categorization stage |  |  |
| --- | --- | --- | --- | --- |
|  |  | Correct rate overall(%) | Trial number/session | Current stage training days |
| mouse1 | 46 | 82.8±19.3 | 589±193 | 11 |
| mouse3 | 37 | 77.2±17.0 | 634±89 | 13 |
| mouse4 | 21 | 88.8±12.8 | 528±110 | 9 |
| mouse6 | 38 | 92.2±6.1 | 672±67 | 6 |
| mouse8 | 35 | 77.2±18.2 | 531±200 | 11 |
| mouse9 | 36 | 88.4±5.7 | 434±176 | 8 |
| o7 | 58 | 69.5±17.5 | 646±94 | 17 |
| rs11 | 23 | 90.9±8.0 | 589±94 | 9 |
| rs3 | 60 | 84.2±13.3 | 620±106 | 4 |
| rs4 | 49 | 90.7±7.9 | 639±55 | 5 |

For each mouse, the overall correct rate and trial number for each session during the 2-tone discrimination stage are shown as mean ± s.d.

**Extended Data Table 2.** Behavior training data for the 1 reversing stimulus stage.

| Mouse name | Training with 1 reversing stimulus |  |  |  |  |  | Current stage training days |
| --- | --- | --- | --- | --- | --- | --- | --- |
|  | Correct rate overall (%) | Non-reversing trial correct rate (%) | Reversing trial correct rate (%) | Correct rate of the first 10 reversing trials (%) | Correct rate of last 10 reversing trials (%) | Trial number/ session |  |
| mouse1 | 83.5±3.1 | 96.1±2.6 | 68.5±5.0 | 43.3±10.5 | 85.3±9.3 | 630±114 | 20 |
| mouse3 | 88.8±2.0 | 98.7±1.0 | 76.0±4.3 | 57.0±11.0 | 88.3±6.2 | 824±124 | 6 |
| mouse4 | 86.4±3.2 | 97.4±1.5 | 71.1±7.4 | 55.3±19.3 | 81.5±7.0 | 459±137 | 5 |
| mouse6 | 86.7±1.4 | 96.6±1.4 | 74.4±1.8 | 47.5±16.4 | 88.9±5.7 | 771±304 | 6 |
| mouse8 | 87.1±2.4 | 96.3±2.9 | 74.9±5.5 | 48.9±11.9 | 92.6±4.0 | 576±110 | 8 |
| mouse9 | 85.6±3.7 | 93.5±3.2 | 76.3±4.9 | 53.4±14.3 | 91.4±6.4 | 590±84 | 9 |
| o7 | 75.8±5.8 | 86.0±6.7 | 56.7±10.4 | 61.3±15.8 | 70.9±14.0 | 724±137 | 26 |
| rs11 | 84.5±3.9 | 96.3±2.6 | 64.3±7.0 | 35.9±11.6 | 84.8±15.4 | 707±102 | 11 |
| rs3 | 86.1±3.3 | 95.3±2.5 | 69.5±7.6 | 54.9±24.5 | 86.1±17.3 | 652±120 | 30 |
| rs4 | 85.3±4.1 | 94.2±7.1 | 71.4±8.9 | 51.0±26.8 | 89.0±7.1 | 739±102 | 16 |
| rs6 | 83.2±2.9 | 94.2±2.8 | 61.8±11.2 | 40.4±18.2 | 83.2±20.0 | 662±76 | 27 |

For each mouse, the overall correct rate, non-reversing trial correct rate, reversing trial correct rate, the correct rate of the first and last 10 reversing trials within each block, and the trial number for each session during the 1 reversing stimulus stage are shown as mean ± s.d.

**Extended Data Table 3.** Behavior training data for the 3 reversing stimulus stage.

| Mouse name | Training with 3 reversing stimulus |  |  |  |  |  | Current stage training days |
| --- | --- | --- | --- | --- | --- | --- | --- |
|  | Correct rate overall (%) | Non-reversing trial correct rate (%) | Reversing trial correct rate (%) | Correct rate of the first 10 reversing trials (%) | Correct rate of last 10 reversing trials (%) | Trial number/session |  |
| mouse1 | 84.3 | 99 | 51.8 | 43.5±10.4 | 86.3±8.4 | 445 | 1 |
| mouse3 | 83.2±0.8 | 95.1±1.1 | 56.7±2.7 | 51.5±13.8 | 85.2±9.2 | 478±52 | 3 |
| mouse4 | 81.9±3.2 | 96.7±2.2 | 52.0±7.9 | 50.6±16.9 | 70.9±16.6 | 542±54 | 4 |
| mouse6 | 79.8±2.7 | 94.4±2.6 | 51.1±9.3 | 47.6±13.7 | 78.5±14.5 | 541±37 | 4 |
| mouse8 | 83.6±4.1 | 94.3±2.6 | 60.7±7.4 | 48.9±13.5 | 86.5±13.2 | 567±170 | 4 |
| mouse9 | 86.6±1.8 | 97.9±0.5 | 58.5±5.7 | 50.1±15.2 | 87.3±9.5 | 605±39 | 2 |
| o7 | 83.2±2.1 | 95.2±1.8 | 61.4±5.0 | 63.8±10.8 | 72.9±14.1 | 659±107 | 15 |
| rs11 | 87.2±0.9 | 98.1±0.8 | 61.2±3.8 | 40.6±14.9 | 81.4±15.7 | 494±128 | 3 |
| rs3 | 85.1±2.8 | 96.5±3.4 | 57.3±5.5 | 55.8±24.3 | 83.3±17.1 | 411±126 | 9 |
| rs4 | 86.1±1.6 | 97.6±1.8 | 58.8±7.2 | 51.8±18.7 | 76.0±15.7 | 648±140 | 20 |
| rs6 | 82.7±1.7 | 92.7±1.8 | 58.9±5.1 | 41.8±18.0 | 79.4±20.5 | 470±111 | 8 |

For each mouse, the overall correct rate, non-reversing trial correct rate, reversing trial correct rate, the correct rate of the first and last 10 reversing trials within each block, and the trial number for each session during the 3 reversing stimulus stage are shown as mean ± s.d.

**Movie S1. Local Ca^2+^ signals from local dendritic tuft regions**

Shown are fluorescence time series at 30 fps from an imaging field from an example behavioral trial.

**Movie S2. Global Ca^2+^ signals from multiple dendritic tuft branches**

Shown are fluorescence time series at 30 fps from the same imaging field as in Movie S1, from an example behavioral trial.

## References

1. Spruston, N. Pyramidal neurons: Dendritic structure and synaptic integration. Nature Reviews Neuroscience 9, 206–221 (2008).

2. Douglas, R. J. & Martin, K. A. C. Neuronal Circuits of the Neocortex. Annual Review of Neuroscience 27, 419–451 (2004).

3. Harris, K. D. & Mrsic-Flogel, T. D. Cortical connectivity and sensory coding. Nature 503, 51–58 (2013).

4. Stuart, G. J. & Spruston, N. Dendritic integration: 60 years of progress. Nat Neurosci 18, 1713–1721 (2015).

5. Dendrites. (Oxford University Press, Oxford, 2016).

6. Major, G., Larkum, M. E. & Schiller, J. Active Properties of Neocortical Pyramidal Neuron Dendrites. Annual Review of Neuroscience 36, 1–24 (2013).

7. Larkum, M. A cellular mechanism for cortical associations: an organizing principle for the cerebral cortex. Trends in Neurosciences 36, 141–151 (2013).

8. Chang, H.-T. Dendritic potential of cortical neurons produced by direct electrical stimulation of the cerebral cortex. Journal of Neurophysiology 14, 1–21 (1951).

9. Larkum, M. E., Zhu, J. J. & Sakmann, B. A new cellular mechanism for coupling inputs arriving at different cortical layers. Nature 398, 338–341 (1999).

10. Magee, J. C. & Johnston, D. A Synaptically Controlled, Associative Signal for Hebbian Plasticity in Hippocampal Neurons. Science 275, 209–213 (1997).

11. Larkum, M. E., Nevian, T., Sandier, M., Polsky, A. & Schiller, J. Synaptic integration in tuft dendrites of layer 5 pyramidal neurons: A new unifying principle. Science 325, 756–760 (2009).

12. Schiller, J., Schiller, Y., Stuart, G. & Sakmann, B. Calcium action potentials restricted to distal apical dendrites of rat neocortical pyramidal neurons. The Journal of Physiology 505, 605– 616 (1997).

13. Losonczy, A., Makara, J. K. & Magee, J. C. Compartmentalized dendritic plasticity and input feature storage in neurons. Nature 452, 436–441 (2008).

14. Helmchen, F., Svoboda, K., Denk, W. & Tank, D. W. In vivo dendritic calcium dynamics in deep-layer cortical pyramidal neurons. Nat. Neurosci. 2, 989–996 (1999).

15. Jia, H., Rochefort, N. L., Chen, X. & Konnerth, A. Dendritic organization of sensory input to cortical neurons in vivo. Nature 464, 1307–1312 (2010).

16. Lavzin, M., Rapoport, S., Polsky, A., Garion, L. & Schiller, J. Nonlinear dendritic processing determines angular tuning of barrel cortex neurons in vivo. Nature 490, 397–401 (2012).

17. Smith, S. L., Smith, I. T., Branco, T. & Häusser, M. Dendritic spikes enhance stimulus selectivity in cortical neurons in vivo. Nature 503, 115–120 (2013).

18. Chen, X., Leischner, U., Rochefort, N. L., Nelken, I. & Konnerth, A. Functional mapping of single spines in cortical neurons in vivo. Nature 475, 501–505 (2011).

19. Beaulieu-Laroche, L., Toloza, E. H. S., Brown, N. J. & Harnett, M. T. Widespread and Highly Correlated Somato-dendritic Activity in Cortical Layer 5 Neurons. Neuron 103, 235-241.e4 (2019).

20. Xu, N., et al. Nonlinear dendritic integration of sensory and motor input during an active sensing task. Nature 492, 247–251 (2012).

21. Kerlin, A., et al. Functional clustering of dendritic activity during decision-making. eLife 8, 1–32 (2019).

22. Francioni, V., Padamsey, Z. & Rochefort, N. L. High and asymmetric somato-dendritic coupling of V1 layer 5 neurons independent of visual stimulation and locomotion. eLife 8, e49145 (2019).

23. Cichon, J. & Gan, W.-B. Branch-specific dendritic Ca2+ spikes cause persistent synaptic plasticity. Nature 520, 180–185 (2015).

24. Otor, Y. et al. Dynamic compartmental computations in tuft dendrites of layer 5 neurons during motor behavior. Science 376, 267–275 (2022).

25. Takahashi, N., Oertner, T. G., Hegemann, P. & Larkum, M. E. Active cortical dendrites modulate perception. Science 354, 1587–1590 (2016).

26. Gold, J. I. & Shadlen, M. N. The Neural Basis of Decision Making. Annual Review of Neuroscience 30, 535–574 (2007).

27. Okazawa, G. & Kiani, R. Neural Mechanisms That Make Perceptual Decisions Flexible. Annu. Rev. Physiol. 85, 191–215 (2023).

28. Hanks, T. D. & Summerfield, C. Perceptual Decision Making in Rodents, Monkeys, and Humans. Neuron 93, 15–31 (2017).

29. Branco, T. & Häusser, M. The single dendritic branch as a fundamental functional unit in the nervous system. Current Opinion in Neurobiology 20, 494–502 (2010).

30. Polsky, A., Mel, B. W. & Schiller, J. Computational subunits in thin dendrites of pyramidal cells. Nat Neurosci 7, 621–627 (2004).

31. Johnston, D., Magee, J. C., Colbert, C. M. & Christie, B. R. Active Properties of Neuronal Dendrites. Annual Review of Neuroscience 19, 165–186 (1996).

32. d’Aquin, S., et al. Compartmentalized dendritic plasticity during associative learning. Science 376, eabf7052 (2022).

33. Mainen, Z. F. & Sejnowski, T. J. Influence of dendritic structure on firing pattern in model neocortical neurons. Nature 382, 363–366 (1996).

34. London, M. & Häusser, M. Dendritic computation. Annu. Rev. Neurosci. 28, 503–532 (2005).

35. Lillicrap, T. P., Santoro, A., Marris, L., Akerman, C. J. & Hinton, G. Backpropagation and the brain. Nat Rev Neurosci 21, 335–346 (2020).

36. Liu, Y., Xin, Y. & Xu, N. A cortical circuit mechanism for structural knowledge-based flexible sensorimotor decision-making. Neuron 109, 2009–2024.e6 (2021).

37. Zhong, L. et al. Causal contributions of parietal cortex to perceptual decision-making during stimulus categorization. Nature Neuroscience 22, 963 (2019).

38. Xin, Y. et al. Sensory-to-Category Transformation via Dynamic Reorganization of Ensemble Structures in Mouse Auditory Cortex. Neuron 103, 909–921.e6 (2019).

39. Hattox, A. M. & Nelson, S. B. Layer V Neurons in Mouse Cortex Projecting to Different Targets Have Distinct Physiological Properties. Journal of Neurophysiology 98, 3330–3340 (2007).

40. Shepherd, G. M. G. Corticostriatal connectivity and its role in disease. Nat Rev Neurosci 14, 278–291 (2013).

41. Chen, X., et al. High-Throughput Mapping of Long-Range Neuronal Projection Using In Situ Sequencing. Cell 179, 772-786.e19 (2019).

42. Jiang, T., Gong, H. & Yuan, J. Whole-brain Optical Imaging: A Powerful Tool for Precise Brain Mapping at the Mesoscopic Level. Neurosci. Bull. (2023) doi:10.1007/s12264-023-01112-y.

43. Gao, L. et al. Single-neuron projectome of mouse prefrontal cortex. Nat Neurosci 25, 515–529 (2022).

44. Gao, L. et al. Single-neuron analysis of dendrites and axons reveals the network organization in mouse prefrontal cortex. Nat Neurosci 26, 1111–1126 (2023).

45. Znamenskiy, P. & Zador, A. M. Corticostriatal neurons in auditory cortex drive decisions during auditory discrimination. Nature 497, 482–485 (2013).

46. Matyas, F. et al. Motor Control by Sensory Cortex. Science 330, 1240–1243 (2010).

47. Tang, L. & Higley, M. J. Layer 5 Circuits in V1 Differentially Control Visuomotor Behavior. Neuron (2019) doi:10.1016/j.neuron.2019.10.014.

48. Takahashi, N. et al. Active dendritic currents gate descending cortical outputs in perception. Nat Neurosci 23, 1277–1285 (2020).

49. Engelhard, B. et al. Specialized coding of sensory, motor and cognitive variables in VTA dopamine neurons. Nature 570, 509–513 (2019).

50. Quian Quiroga, R. & Panzeri, S. Extracting information from neuronal populations: information theory and decoding approaches. Nat Rev Neurosci 10, 173–185 (2009).

51. Domnisoru, C., Kinkhabwala, A. A. & Tank, D. W. Membrane potential dynamics of grid cells. Nature 495, 199–204 (2013).

52. Dayan, P. & Abbott, L. Theoretical Neuroscience: Computational And Mathematical Modeling of Neural Systems. (Massachusetts Institute of Technology Press, 2005).

53. Murayama, M. et al. Dendritic encoding of sensory stimuli controlled by deep cortical interneurons. Nature 457, 1137–1141 (2009).

54. Ma, Y., Hu, H., Berrebi, A. S., Mathers, P. H. & Agmon, A. Distinct Subtypes of Somatostatin-Containing Neocortical Interneurons Revealed in Transgenic Mice. J. Neurosci. 26, 5069–5082 (2006).

55. Higley, M. J. Localized GABAergic inhibition of dendritic Ca2+ signalling. Nat Rev Neurosci 15, 567–572 (2014).

56. Chiu, C. Q. et al. Compartmentalization of GABAergic Inhibition by Dendritic Spines. Science 340, 759–762 (2013).

57. Tremblay, R., Lee, S. & Rudy, B. GABAergic Interneurons in the Neocortex: From Cellular Properties to Circuits. Neuron 91, 260–292 (2016).

58. Branco, T. & Häusser, M. Synaptic Integration Gradients in Single Cortical Pyramidal Cell Dendrites. Neuron 69, 885–892 (2011).

59. Losonczy, A. & Magee, J. C. Integrative Properties of Radial Oblique Dendrites in Hippocampal CA1 Pyramidal Neurons. Neuron 50, 291–307 (2006).

60. Grienberger, C., Chen, X. & Konnerth, A. NMDA Receptor-Dependent Multidendrite Ca 2+ Spikes Required for Hippocampal Burst Firing In Vivo. Neuron 81, 1274–1281 (2014).

61. Fişek, M. et al. Cortico-cortical feedback engages active dendrites in visual cortex. Nature 617, 769–776 (2023).

62. Voigts, J. & Harnett, M. T. Somatic and Dendritic Encoding of Spatial Variables in Retrosplenial Cortex Differs during 2D Navigation. Neuron 105, 237–245.e4 (2020).

63. Economo, M. N. et al. Distinct descending motor cortex pathways and their roles in movement. Nature 563, 79 (2018).

64. Constantinople, C. M. & Bruno, R. M. Deep Cortical Layers are Activated Directly by Thalamus. Science 340, 1591–1594 (2013).

65. Harris, K. D. & Shepherd, G. M. G. The neocortical circuit: themes and variations. Nat Neurosci 18, 170–181 (2015).

66. Zingg, B. et al. Neural Networks of the Mouse Neocortex. Cell 156, 1096–1111 (2014).

67. Winkowski, D. E. et al. Orbitofrontal Cortex Neurons Respond to Sound and Activate Primary Auditory Cortex Neurons. Cereb Cortex 28, 868–879 (2018).

68. Schneider, D. M., Nelson, A. & Mooney, R. A synaptic and circuit basis for corollary discharge in the auditory cortex. Nature 513, 189–194 (2014).

69. Lefort, S., Tomm, C., Floyd Sarria, J.-C. & Petersen, C. C. H. The Excitatory Neuronal Network of the C2 Barrel Column in Mouse Primary Somatosensory Cortex. Neuron 61, 301– 316 (2009).

70. Jiang, X. et al. Principles of connectivity among morphologically defined cell types in adult neocortex. Science 350, aac9462–aac9462 (2015).

71. Fu, Y. et al. A Cortical Circuit for Gain Control by Behavioral State. Cell 156, 1139–1152 (2014).

72. Lee, S., Kruglikov, I., Huang, Z. J., Fishell, G. & Rudy, B. A disinhibitory circuit mediates motor integration in the somatosensory cortex. Nat Neurosci 16, 1662–1670 (2013).

73. Poort, J. et al. Learning and attention increase visual response selectivity through distinct mechanisms. Neuron 110, 686–697.e6 (2022).

74. Pi, H.-J. et al. Cortical interneurons that specialize in disinhibitory control. Nature 503, 521–524 (2013).

75. Guo, Z. V. et al. Procedures for behavioral experiments in head-fixed mice. PLoS ONE 9, (2014).

76. Pachitariu, M., et al. Suite2p: beyond 10,000 neurons with standard two-photon microscopy. bioRxiv 061507 (2016) doi:10.1101/061507.

77. Kraskov, A., Stögbauer, H. & Grassberger, P. Estimating mutual information. Phys. Rev. E 69, 066138 (2004).

78. Ross, B. C. Mutual Information between Discrete and Continuous Data Sets. PLOS ONE 9, e87357 (2014).

79. Gong, H., et al. Continuously tracing brain-wide long-distance axonal projections in mice at a one-micron voxel resolution. NeuroImage 74, 87–98 (2013).

